# Recording from the same neuron with high-density CMOS probes and patch-clamp: a ground-truth dataset and an experiment in collaboration

**DOI:** 10.1101/370080

**Authors:** André Marques-Smith, Joana P. Neto, Gonçalo Lopes, Joana Nogueira, Lorenza Calcaterra, João Frazão, Danbee Kim, Matthew G. Phillips, George Dimitriadis, Adam R. Kampff

## Abstract

We built a rig to perform patch-clamp and extracellular recordings from the same neuron *in vivo.* In this setup, the axes of two micromanipulators are precisely aligned and their relative position tracked in real-time, allowing us to accurately target patch-clamp recordings to neurons near an extracellular probe. We used this setup to generate a publicly-available dataset where a cortical neuron’s spiking activity is recorded in patch-clamp next to a dense CMOS Neuropixels probe. “Ground-truth” datasets of this kind are rare but valuable to the neuroscience community, as they power the development and improvement of spike-sorting and analysis algorithms, tethering them to empirical observations. In this article, we describe our approach and report exploratory and descriptive analysis on the resulting dataset. We study the detectability of patch-clamp spikes on the extracellular probe, within-unit reliability of spike features and spatiotemporal dynamics of the action potential waveform. We open discussion and collaboration on this dataset through an online repository, with a view to producing follow-up publications.

**Prologue:** Our efforts to record from the same neuron in vivo using patch-clamp and dense extracellular probes have resulted in three outputs: a publicly-available dataset (http://bit.ly/paired_recs), a manuscript, and a code repository (http://bit.ly/paired_git). Together, these three components form the publication arising from the experiments we have performed. The role of the dataset is to be downloaded and re-used. The role of the manuscript is to describe the experimental methods through which we acquired the dataset, explain it and showcase which types of questions it can be used to address. The repository has two roles: first, promoting reproducibility and error correction. By making our analysis and figure-generation code freely-available, we wish to make our analysis procedures clear and enable the reader to reproduce our results from the raw data, alerting us to any potential mistakes. Second, the repository will form a living, dynamic and interactive component of the publication: a forum for open collaboration on this dataset. Any interested scientists can contribute to it, joining us in detailed exploration of these recordings with a view to producing follow-up publications in which they will be credited for their input.

Why did we opt to publish this way? The first reason is that the very nature of the project we here describe – recording the same neuron with patch-clamp and extracellular probes – invites an open science and open source approach. This is because the primary use of this type of “ground truth” validation data is to aid the development of new sorting and analysis algorithms, as well as to benchmark and improve existing ones. The second reason is that despite being conceptually very simple, this project generated a large and complex dataset that can be tackled in many ways and used to address different types of question. Some of these questions are beyond the reach of our analytical expertise; others lie even beyond the scope of our scientific imagination. By releasing the dataset and providing a repository for scientific discussion and collaboration, we aim to maximise its scientific return to the community. Instead of having each interested research group work in isolation, we hope that by encouraging collaboration and discussion between peers we can foster synergy between them that will lead to work of greater scientific value.

Although datasets like ours are exquisitely suited for such an approach, we believe this publication strategy needs to become more widely adopted in neuroscience. We were pleased to note recent publications spontaneously and independently using similar approaches^1-4^, in what may well be evidence of convergent thinking. Perhaps the time has come for new publication and collaboration paradigms. We will elaborate on this subject during the Epilogue. For now, let us get back to electrophysiological recordings, before we begin an experiment on scientific collaboration.

## Introduction

### Extracellular recordings

Understanding how the brain works requires seeing the forest *and* the trees: we must track large populations of neurons, but also resolve their activity as units^5,6^. As a method, extracellular recordings have come a long way towards achieving this goal, progressing through technological advances from recording single neurons in the 1950s^7-9^ to several hundreds in the 2010s^10-12^. However, with each leap in technology come new questions about the nature of the signal being recorded and requirements for new analysis methods to interpret the ever-growing datasets. Extracellular recordings offer unparalleled access to large populations of neurons deep in the brain with sub-millisecond temporal resolution. However, each electrode can detect the spiking activity of tens to hundreds of neurons nearby, which poses a formidable analytical challenge: how to resolve this chaos of activity into individual units? For this reason, analytical innovations have been just as important as new recording technologies in increasing the power of extracellular recordings^13^.

Complementary metal-oxide-semiconductor (CMOS) probes are the latest advance in extracellular recording technology^12^. CMOS probes exploit innovations in microfabrication techniques that ultimately enable scientists to record from hundreds of channels (384 to 1,440), densely packed (100-170 sites/mm) along a 5-10 mm shank^12^. These devices enable access to hundreds of neurons distributed across multiple brain regions, densely sampling the extracellular field; this means that each neuron is detected on multiple channels of a probe. This additional resolution in sampling is expected to aid analysis and is currently being used by novel algorithms^14-17^.

### “Ground truth” data

Datasets where one knows precisely when a neuron in the vicinity of an extracellular probe fired an action potential (commonly referred to as “ground-truth” ^1^) are necessary for validating the performance of new recording technology and benchmarking analysis approaches. They have also been essential for advancing our understanding of the nature of the extracellular signal and how it corresponds to intracellular recordings^18^. Finally, they can provide empirical answers to matters of technological design: is it more useful to optimise electrode arrangement for drift correction, or to position channels strategically in the hope of isolating more units? These datasets have been acquired for tetrodes/single-wire electrodes^18–21^ or in slice preparations^22^, where background activity is greatly reduced. We recently added to this literature by publishing and sharing a ground-truth dataset from silicon polytrodes^23^, but these devices have significantly different channel count and arrangement from the new CMOS probes^10,11^. Ground truth datasets from tetrodes in the hippocampus^18^ and polytrodes in cortex^23^ have proved invaluable for constraining, benchmarking and improving spike sorting algorithms^15,24–27^. Given the potential of CMOS probes for becoming standardised tools adopted worldwide^12^, it would be of great interest to produce a ground-truth dataset for these new devices that can be used to develop common analysis tools and standards. That is what we did here, using our previously published method for efficiently targeting two different electrophysiological recording instruments to the same neuron *in vivo*^23^. The dataset is now available online (http://bit.ly/paired_recs) and the reader can collaborate with others and ourselves in its analysis (http://bit.ly/paired_git).

## Materials and Methods

### Dual-Recording Rig Design and Alignment

For this project, we adapted the design of our previous dual-recording rig^23^. Our implementation of a dual-recording rig requires two aligned multi-axis micro-manipulators (*Scientifica Patchstar*), a long working distance custom-built optical microscope (*Optomechanics: Thorlabs; Objective: Mitutoyo 378 series 10x; Camera: PointGrey Flea3 USB*) to align the extracellular probe and patch pipette tip, a macro-zoom lens (*Edmund Optics 3.3X Macro Zoom Lens, coupled to a second PointGrey Flea3 USB camera*) to guide probe and patch pipette insertion, software (*NeuroGEARS Bonsai*^28^) to monitor probe and patch pipette position and calculate distance between pipette tip and a given coordinate on the probe in real time, a stereotaxic frame for rat head fixation (*Kopf Model 962*), a computer, and electrophysiology acquisition hardware (described below in “*Experiments”* section). The air table on which the stereotaxic frame was mounted defined the common X-, Y- and Z- axes to which manipulators were aligned: X is parallel to the anatomical medio-lateral axis (ML), Y to the anterior-posterior axis (AP) and Z to the dorso-ventral axis (DV). The two Patchstar manipulators were mounted on opposite sides of the stereotaxic frame along the X-axis. They were held at an angle: the probe manipulator at 64° from X on the XZ plane, and the patch manipulator at 62° on the same plane.

#### Mechanical alignment

After assembling the rig, we ensured that the axes of both manipulators were parallel using a mechanical alignment procedure. We “squared” manipulators with the air table surface using a digital machinist’s dial (*RS Pro Fine Reading Indicator*) mounted on the electrode holder. We placed the dial tip in contact with a planar surface of the air table and moved it along this surface (see ref. 23 for a video of this procedure). The dial is sensitive to changes as small as 1 μm; changes > 50μm for the full range of travel of the manipulator were corrected by loosening or tightening manipulator mounting screws or tapping the manipulator with a soft surface hammer until such differentials were minimised. We performed this procedure for X-, Y- and Z-axes.

#### Optical Alignment

The numerical aperture of the alignment microscope objective (0.28) has a theoretical resolution limit of 1 μm in the X- and Y-axes and 10 μm in Z. Before each experiment, we mounted a model rat skull (with bregma, lambda and a craniotomy) on the stereotaxic frame. The alignment microscope’s objective was focused on a point a few hundred to 1,000 μm above bregma. We brought the probe and patch pipette tips to this point, illuminating them obliquely in order to acquire images of both with sufficient contrast. We aligned probe and patch pipette tips visually at the centre of the image (indicated by an overlay crosshair), and reset their XYZ coordinates to zero on the manipulator position monitoring software (Scientifica Linlab 2.0), a procedure we refer to as “zeroing”. We previously verified the repeatability of zeroing by manually moving the tip of the pipette from outside the field of view to the focal plane and image centre and recording the manipulator coordinates, having found a 0.5 ± 0.5 μm reliability in XY and 2.6 ± 1.7 μm in Z^23^. We then moved the probe to a different position in space. The microscope was moved and refocused to re-centre the probe tip at the crosshair. Next, without moving the microscope, we moved the pipette tip to the same XYZ coordinates as the probe tip. If there was no misalignment between manipulators, the pipette tip should arrive at the centre of the image crosshair, that is, the same position as the probe tip. In the event of misalignment, the amount of re-positioning required (in X, Y and Z) to bring the pipette tip to the probe tip provides an accurate measure of residual axis misalignment. We performed this procedure sequentially to 15 different locations in XYZ spanning a 5,000 μm x 5,000 μm x 5,000 μm volume in 1000 μm steps recording every time the cumulative displacement in X, Y and Z. The average distance error recorded in this volume was 9.9 ± 6.2 μm (n = 30 measures).

#### Software alignment

Taking the probe manipulator as reference, we can use the position errors measured at several different locations to estimate the coordinate transformation that best compensates for any residual misalignment of the patch pipette manipulator. We used multivariate linear regression to compute a transformation matrix for XYZ and then used the constants derived to transform patch manipulator coordinates in real time using Bonsai, an open-source reactive visual programming framework (Lopes et al., 2015; freely available for download at http://bonsai-rx.org/). This procedure allowed us to reduce residual misalignment to 6.8 ± 3.6 μm. The Bonsai workflow we used is available for download at the sc.io repository.

#### Bonsai-guided targeting of patch-clamp recordings

Having aligned probe and patch pipette manipulators with sufficient accuracy, we required a way to calculate XYZ coordinates for a patch pipette entry point into the brain that would allow us, following a straight line path through the brain, to reach a point in space sufficiently close to one of the probe’s channels. Our approach was the following: first, we reset probe and patch pipette tip coordinates to zero, at a fixed point located a few hundred microns over bregma (Figure 1A). Second, we guided the probe to the target craniotomy location and lowered it slowly until we could detect a dimple on the surface of the brain (Figure 1B). Third, we recorded the XYZ coordinates for this point (probe entry, point A Figure 1D), zeroed the virtual approach axis (XZ) to monitor how deeply the probe was implanted (range from 2500 μm to 3800 μm for different experiments), inserted the probe into the brain at an approach angle of 64° and recorded the final coordinates for the probe tip (point B Figure 1D). This left us with a line segment AB defined by two sets of XYZ coordinates: A (probe entry) and B (probe tip), as depicted in Figure 1D. This allowed us to derive a “probe line equation” of the form Z = mX + b where m is the slope (64°) and b is the intercept. Fourth, every time we “hunted” for a cell, we picked a point T (target) located along the probe line segment. Knowing this point and our patch manipulator approach angle (62°), we could now define a “patch line equation” for the line that passes through this point at 62° to the horizontal (segment CT in Figure 1D). By fixing an arbitrary patch pipette Z coordinate to a point above the brain surface (e.g. points C or C’, Figure 1D), we could solve the patch line equation for X and thus obtain our X axis (ML) entry point. At this stage we are therefore in possession of the three required coordinates for a patch pipette entry point through which we may ultimately reach point T on the probe. To reiterate, the Z coordinate is defined arbitrarily and based on convenience of movement; picking a Z coordinate very far above the brain surface will require greater movement along the X axis (see Figure 1D, point C versus C’), which may be impractical or outside the travel range of the manipulator. The X coordinate is calculated based on the arbitrarily-defined Z. Finally, the Y coordinate is the same as for the probe. In possession of 3 Cartesian coordinates for an entry point C (Figure 1D) we are therefore ready to slowly advance into the brain along the approach line CT (Figure 1D) that will lead us to a region in space close to the extracellular probe, hoping to “collide” with and obtain patch-clamp recordings from neurons in this path. We implemented these calculations and variables in a Bonsai workflow, enabling us to a) obtain patch entry point coordinates “on the fly” and b) estimate in real time the current Euclidean distance between the patch pipette tip and the closest point on the probe. The latter allowed us to both avoid colliding the patch pipette with the probe and – crucially – estimate the distance between a recorded neuron and its closest point in the probe with accuracy, obviating the need for (potentially loss-prone) post-hoc histology of patch-clamp recorded neurons. The approach we just described was tested “in air” prior to beginning recordings on each experiment day. Briefly, a virtual probe insertion was performed using the same coordinates as chosen for the experiment. We calculated the patch entry point, focused the microscope on the target point of the probe and moved the patch pipette along the approach line, visually confirming that its tip touched the target point. If there was a displacement of > 10 μm between pipette tip and target, we repeated optical and software alignment procedures from the beginning.

**Figure 1.**
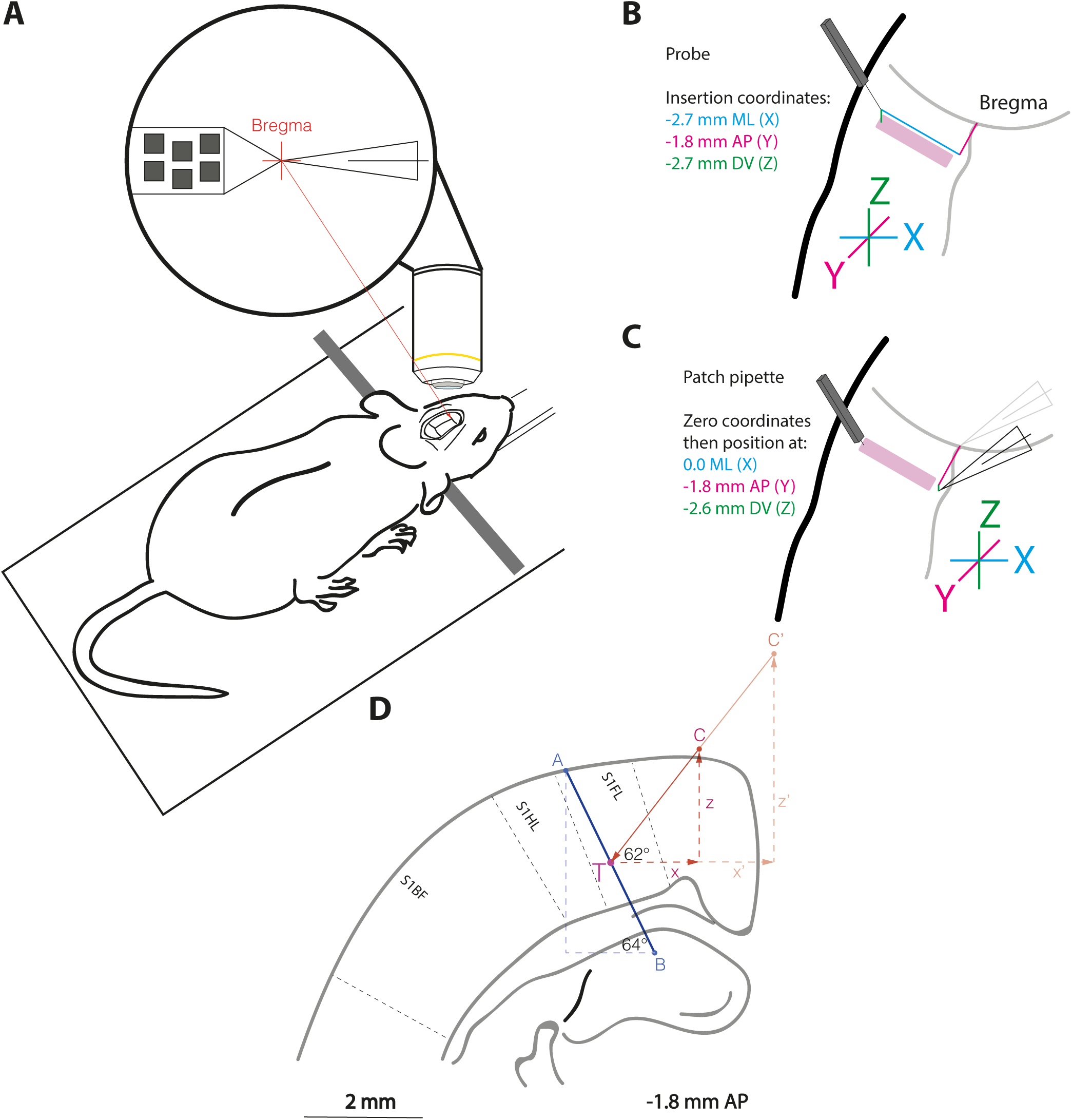
Bonsai-guided targeting for paired recordings. A. At the beginning of each experiment, the alignment microscope objective was focused on and fixed at a reference point a few hundred microns above bregma (crosshair). The probe and patch pipette tips were brought to this point and zeroed. B. We then guided the probe to the insertion point, noted coordinates and implanted the probe deep enough to record from all cortical layers. C. For every patch-clamp recording, the pipette was brought back to the alignment point depicted in A and its coordinates reset to 0. It was then moved to the target Y (AP) coordinate and a fixed Z (DV) position. D. We then picked a target location on the probe (T), defining a point in XYZ to target with patch-clamp. Depending on the fixed Z position decided upon, this returned a different XY entry point (points C, C’). The pipette tip was positioned at the entry point, switched to approach mode, and advanced slowly into the brain at a 62 degree angle.

### Experiments

#### Surgery

All animal experiments were approved by the University College London local ethical review committee and conducted in accordance with Home Office personal and project (I6A5C9913; 70/8116) licenses, under the UK Animals (Scientific Procedures) 1986 Act.

We used Lister-Hooded rats of both sexes, aged between 6 weeks and 8 months (weight 300-700 g). Rats were anaesthetized with a single injection of urethane (1.4-1.8 g/kg intraperitoneal), which was followed by a subcutaneous injection of temgesic (20 μg/kg) and rimadyl (5 mg/kg) and intra-muscular injection of Atropine methyl-nitrate (0.05 mg/kg), for suppression of mucus secretion. Depth of anesthesia was monitored by paw and tail-pinch and after 2 hours if no pain reflex was observed, surgical procedures were initiated. Rats were mounted on a stereotaxic frame and their temperature monitored rectally and maintained at 37.5 °C by a homeothermic blanket. Lidocaine was injected subcutaneously along the midline of the scalp. We performed an incision to expose the skull above the targeted brain region(s). One or two line-shaped craniotomies (1mm along the AP and 2-3 mm along the ML axes) were performed. An incision was performed with a thin scalpel blade or bent 29G needle on the underlying dura along its full ML extent, taking care to minimize the area of brain surface exposed and not damage it. Craniotomy coordinates are detailed for each experiment on the accompanying “Data Summary” spreadsheet. Two reference electrodes (Ag-AgCl wires from Science Products, model E-255) were implanted opposite each other, under the most posterior section of the skin incision just above the neck.

#### Extracellular Probes

All experiments were performed with Neuropixels Phase3A Option 1 Probes (IMEC). This probe model has 384 channels, each with an area of 144 μm^2^, arranged in a chessboard pattern along 192 rows and 4 columns^10^. Columns are spaced 21 μm and rows 20 μm apart (see Figure 2 below; also https://github.com/cortex-lab/neuropixels/wiki). Probe shank dimensions are 5 mm length by a 70×20 μm cross-section. We were unable to measure probe channel impedance but Jun, Steinmetz and colleagues have previously documented this value to be 149 ± 6 KΩ (Mean ± SD)^10^. In Neuropixels probes, the continuous data stream from each channel is split into action potential (AP, 0.3-10,000 kHz) and local field potential bands (LFP, 0.5-1,000 Hz), which are amplified and digitized separately (AP 30 kHz, LFP 2.5 kHz). Digitization was performed at 10 bits, under a gain of 500, yielding a resolution of 2.34 μV per bit. We acquired data using SpikeGLX open-source software (https://github.com/billkarsh/SpikeGLX).

**Figure 2.**
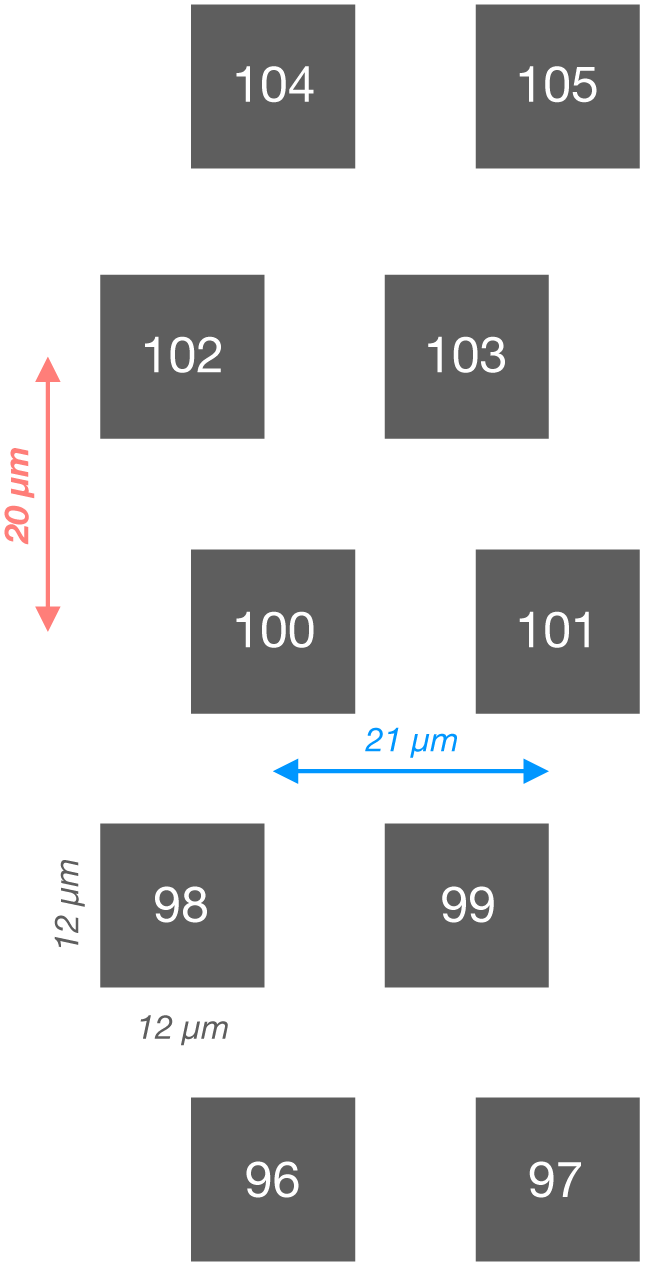
Neuropixels Option 1 probe layout. Example channel numbers in white, embedded within channel (grey). High channel numbers are towards brain surface.

#### Patch-Clamp recordings

We used filamented borosilicate glass capillaries (OD 1.5mm, ID 0.86 mm, 10 cm length; WPI) and pulled patch pipettes from them using a Narishige PC-10 vertical puller, configuring a pulling program to yield pipette tips with resistance between 7 and 9 MΩ. Patch pipettes were filled with intracellular solution containing (in mM): 130 KGluconate, 5 KCl, 10 HEPES, 2 MgCl_2_, 10 Sodium Phosphocreatine, 2 Na_2_-ATP and 0.4 Na-GTP. For patch-clamp recordings, data was acquired using an Axon Instruments Multiclamp 700B amplifier and a National Instruments board, at a sampling rate of 50.023 kHz. Please see accompanying Data Summary spreadsheet to verify sampling rate for each cell.

We used WinWCP 5.3.4 software for acquisition (developed by John Dempster, freely-available at http://spider.science.strath.ac.uk/sipbs/software_ses.htm).

#### Dual Recordings with Extracellular Probes and Patch-Clamp

At the beginning of an experiment, after all alignment steps detailed above, extracellular probe position was zeroed at the center of a virtual crosshair positioned ~1mm over the rat’s bregma (Figure 1). The extracellular probe was then guided to the craniotomy site and lowered until a dimple was observed at the brain surface (see *Bonsai-guided targeting of patch-clamp recordings*). The probe was then inserted at a constant velocity of 5 μm/s. We allowed 30 minutes for brain tissue around the probe to settle before attempting patch-clamp recordings. At this point, a patch pipette was filled with intracellular solution, brought to the center of the overlay crosshair and zeroed (Figure 1C). The patch pipette was then guided to the calculated entry point coordinates (see *Bonsai-guided targeting of patch-clamp recordings*), at which point we followed a protocol for *in vivo* patch-clamp recordings in rodents^29^. High positive pressure (30-60 mmHg; *DPM1B Pneumatic Transducer Tester; Fluke Biomedical*) was applied before entering the pia and pipette resistance measured in voltage-clamp mode, using 25 mV steps at 20 Hz. Usually, we observed a transient increase in pipette resistance of up to 100%. We advanced the pipette tip steadily and monitored the distance to the target point on the probe. Once this fell below 200 μm, positive pressure was decreased to 10-20 mmHg, and the pipette advanced slowly at 1-2 μm per second. We monitored the test pulse for increases in pipette resistance of ~50% and the appearance of a “strike” pulse^29^, as well as any spike-like waveforms appearing on the recording trace. When a combination of these signs was detected, we released pressure and applied slight suction to attempt to obtain a seal on the putative neuron’s membrane. In cases where a seal resistance of > 1 giga-Ω was obtained, we attempted to go into whole-cell mode by applying short, sharp suction to rupture the cell membrane. In situations where resistance did not reach giga-Ω level, we remained and recorded in cell-attached mode, either in voltage-clamp mode (n = 30 cells), with holding voltage set to yield a current of ~0 pA, or current-clamp mode (n = 8 cells), without injecting current. The remaining 5 cells in the dataset were recorded in whole-cell configuration. Prior to beginning a dual recording, we monitored patch-clamp activity for spikes. For cells without spontaneous spiking activity, we attempted to induce action potentials by injecting current (whole-cell or cell-attached current-clamp) or setting voltage-clamp holding potential to +10 to +50 mV^30^. If the cell was still “silent”, we discarded it and attempted to record from another neuron. Once we observed stable activity with action potentials in patch-clamp, we started a dual recording by initializing a protocol in WinWCP. SpikeGLX extracellular recordings were triggered by the onset of a TTL pulse, commanded by WinWCP and delivered through the National Instruments board to the Neuropixels Sync Channel 0. To ensure that extracellular and patch signals were temporally aligned, TTL pulses were delivered throughout the recording. For each new recording, the pipette was removed from the brain through the approach track and a new pipette filled and mounted. Every new pipette was zeroed as per the procedure described above. We tested in air whether manipulation of the holder during pipette replacement disrupted alignment and we were satisfied that it did not (data not shown).

### Processing and Analysis

#### Inclusion criteria

Unless otherwise specified in particular sections of this article, out of the total of 43 paired recordings obtained, a subsample of 21 was selected for further analysis on the basis of 3 criteria being met: 1) detection of > 200 spikes on the patch-clamp recording; 2) Patch spike-triggered average of the extracellular voltage (PSTA) revealing a waveform with peak-peak amplitude > 10 μV; 3) PSTA revealing a canonical spike waveform, defined as i) a sharp negative peak followed by a positive peak; or ii) a positive peak, followed by at least one negative peak of greater amplitude. Sixteen cells showed peak-peak amplitudes <10 μV; 2 cells fired less than 200 spikes over the whole recording session; 4 cells showed non-canonical spike waveforms. The spike waveforms for excluded and included cells with peak-peak amplitudes > 10 μV are plotted in Supplement to Figure 4-1 A. All recordings are available for download; exclusions related only to analysis subsequent to Figure 4.

#### Channel with highest peak-peak vs Channel estimated to be closest to soma

We used the coordinates returned by our Bonsai workflow for probe and patch pipette tip positions along with the Neuropixels Option 1 probe map to estimate which was the extracellular channel closest to the soma/patch pipette tip. This allowed real-time monitoring of activity in the range of channels most likely to be picking up activity from a dual-recorded cell. Moreover, given that the amplitude of extracellular action potentials is highest in the perisomatic region, this also allowed us to test *in vivo* and inside the brain the accuracy of our alignment and positioning system. Figure 2 is a schematic of the Neuropixels Option 1 probe geometry. In our calculations we accounted for a distance of 137 μm between the sharp tip of the Option 1 probe and the deepest channel (Jun et al., 2017; https://github.com/cortex-lab/neuropixels/wiki). Our estimate for channel predicted to be closest to the soma/pipette tip was calculated thus:

*Predicted channel = 2 * ((1 – fraction_channel) * rows_inside) – 2*,

Where:

*fraction_channel = targeted channel depth / total probe insertion length*
*rows_inside = total probe insertion length / row spacing*
*row spacing = 20 μm*
*targeted channel depth: see section “Bonsai-guided targeting of patch-clamp recordings”*

The difference between predicted channel and channel with the highest peak-peak amplitude is reported in Data Summart. For 70% of recordings with clear EAP waveforms (15/21), our estimate was off by 5 channels or less. Given the geometry arrangement of Neuropixels Option 1 probes (Figure 2), and depending on which column we were closest to within a row, this corresponds to a displacement in Euclidean distance ranging from 45.2 μm (channel at the beginning of a row, for example channel 100 to 105) to 63.6 μm (channel at the end of a row, for example channel 97 to 102). This is a considerable increase compared to the precision of 6.8 ± 3.6 μm obtained in air after software alignment, but it is to be expected, given differences between moving a pipette tip or probe in air versus inside brain tissue, where they may encounter obstacles of varying rigidity such as blood vessels and thicker neurites. Moreover, putting this measure in the context of somatic diameter, if we assume this to be approximately 20 μm for a rat layer 5 cortical neuron^31^, this suggests our estimated versus real pipette tip position inside the brain *in vivo* was predominantly accurate within a range of 2-3 cell bodies. In every analysis using “nearest channel to the soma”, for the minority of situations where there was a difference between channel with highest extracellular amplitude and the one estimated to be nearest the soma, we use the channel with highest extracellular amplitude. Because this reading and spatial estimates were so close, throughout the paper we use the terms “nearest channel to the soma” and “channel with the highest extracellular amplitude” interchangeably.

#### Synchronization

To make sure our dual modality recordings were temporally aligned, we programmed digital pulses, which were recorded simultaneously by the extracellular and patch-clamp acquisition systems. Digital pulses comprised 3 types, distinguished by their time of onset and duration: a “trial initiation pulse” delivered every 30 seconds, a series of “ongoing trial pulses” delivered every second and a “trial conclusion pulse” delivered every 30 seconds. Trial initiation pulses had a duration of 10 ms, whereas ongoing trial pulses lasted 20 ms and trial conclusion pulses 100 ms. We verified extracellular-patch temporal alignment by comparing trial duration for each trial recorded; no misalignments were observed and therefore trials were concatenated for analysis. Since extracellular and patch-clamp recordings were acquired at distinct sampling rates, to relate events in these two streams we used a conversion factor. This was calculated by dividing the sample length of the two streams, obtaining a constant which was then multiplied/divided by the sample number of a particular extracellular/patch-clamp event.

#### Preprocessing

The signal recorded with Neuropixels Phase 3A probes has an offset that varies per channel. To remove this, we subtracted from each channel the median of its voltage. After that, extracellular recordings were filtered using a sixth-order Butterworth high-pass filter at the cutoff frequency of 200 Hz, run in forward and backward mode. Finally, we performed common-average referencing by subtracting from each channel the median voltage over the whole probe at each time-point. The latter two steps were performed only for our own analysis in this article; the data we have shared publicly has only undergone offset subtraction. Patch-clamp data were preprocessed only for spike detection. For this analysis step, we filtered the recording using a sixth-order Butterworth high-pass filter at the cutoff frequency of 100 Hz, run in forward and backward mode.

#### Spike Detection

All code used for analysis and figure production is freely available and commented at the sc.io repository (http://bit.ly/paired_git). Spikes in patch-clamp recordings were detected after high-pass filtering (cutoff frequency 100 Hz) and median-subtracting the data, as crossings of a threshold defined as 7 times the standard deviation of current or voltage across the whole recording. Extracellular spikes were detected as threshold crossings for 7 times the median absolute deviation of the signal within a particular channel. Spike features were thus defined: peak-peak amplitude – voltage at the positive peak minus voltage at the negative peak; half-width – duration at the half-amplitude of the negative peak; duration – time elapsed between the positive and negative peaks; peak-peak ratio – absolute value of the negative peak amplitude divided by positive peak amplitude; symmetry – ratio between time elapse from baseline to negative peak and time elapsed from negative peak back to baseline value.

## Results

Our goal was to record from the same neuron *in vivo* using high-density CMOS probes and patch-clamp. To achieve this, we built a dual-recording rig where the axes of two micromanipulators are precisely aligned and their positions are tracked in real-time by Bonsai software, enabling us to accurately estimate the distance between a patch-clamp recorded neuron and the closest channel on the probe (ref. 23; see Materials and Methods). Our overall aim was to generate and share a ground-truth dataset for Neuropixels, which are part of a new generation of extracellular probes with high channel count and density^10,11^.

### Paired recording: a walk-through

Figure 3A depicts a 100-ms segment of a paired recording where we recorded in voltage-clamp cell-attached mode from a neuron (c46) that was 65 μm away from the probe. Near-simultaneous spikes appear on the patch-clamp trace (Fig. 3A, black) and the nearest channel of the Neuropixels probe (channel 181, Fig. 3A, blue). However, single channels on extracellular probes detect the activity of tens of nearby neurons – could this apparent correspondence in cell-attached and extracellular spike times be a mere coincidence? Randomly sampling 500 spikes from this cell-attached recording and plotting the corresponding time-windows for the nearest Neuropixels channel suggests not; a clear time-locked extracellular spike waveform is seen on the extracellular recording for each and every cell-attached spike (Fig. 3B). Confident that we can track the spiking activity of the same neuron in patch-clamp and extracellular recordings, let us look now to panels 3C-E. While we know that one channel can pick up spikes from several neurons beyond the dual-recorded one (Fig. 3A blue), the converse is also true: spikes from the same neuron can be recorded on multiple channels. Figure 3C depicts the same 100-ms segment of extracellular activity shown in 3A for each of the 30 channels nearest to the probe; however, due to background activity and electrical noise, the individual spike waveforms can sometimes be hard to discern. We therefore computed the patch spike-triggered average (PSTA) by averaging extracellular activity in every channel of the Neuropixel probe in the 4-ms time window surrounding each spike recorded in patch-clamp (Fig. 3D-E). For c46, averaging activity in time windows around 6,803 spikes revealed a variety of distinct extracellular spike waveforms, spread along 30 channels of the Neuropixels probe (Fig. 3D-E). Waveforms obtained for each channel show distinct features in time (Fig. 3D) and space (Fig. 3E). Though we did not recover morphological reconstructions in this study, for every experiment we sought to keep probe insertion angle parallel to the dorso-ventral axis defined by the apical dendrites of cortical pyramidal neurons. As such, channels in rows above the nearest channel are likely sampling voltage near the cell-attached neuron’s apical dendrites, whereas those in rows below potentially capture electric fields near the basal aspect of the dendritic tree.

**Figure 3.**
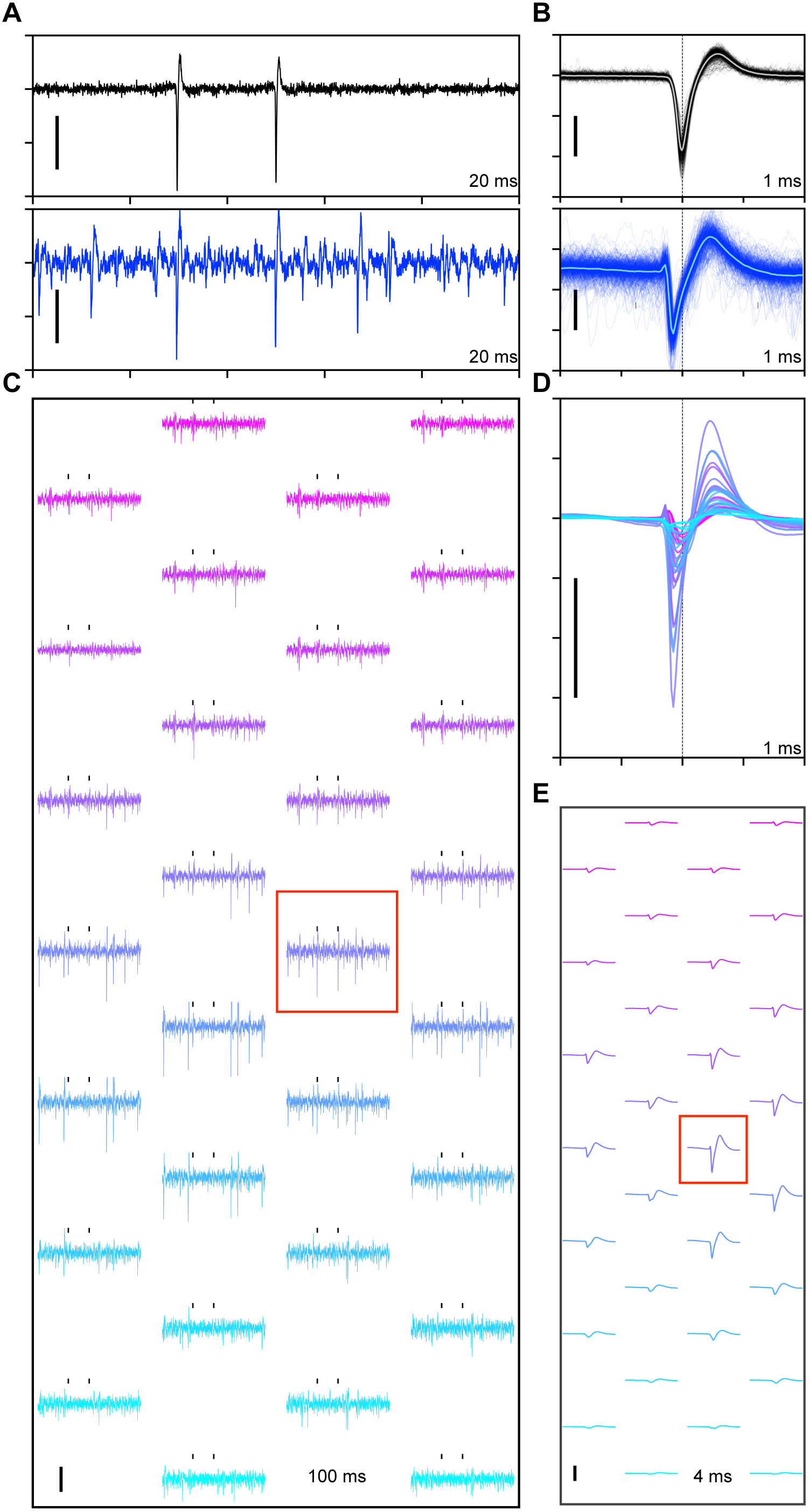
Paired cell-attached and extra-cellular recordings from the same neuron. **A.** A segment of 100ms of cell-attached (top, black) and extra-cellular (bottom, blue) recordings (cell c46). Extra-cellular trace was sampled from the channel closest to the cell body (181). Scale bars for this and subsequent plots are 100 pA (cell-attached) or 100 μV (extracellular). **B.** Sample of 500 cell-attached spikes (top, black) and corresponding extra-cellular 4-ms recording windows from channel 181 (bottom, blue). Trace averages overlaid in lighter tones. **C.** The same temporal segment as in **A** for the 30 channels closest to the cell soma. Channel 181 is indicated by red box. Cell-attached spike times indicated by black ticks. **D.** Patch spike triggered-average (PSTA) of extra-cellular voltage windows for 30 channels closest to the cell body aligned by patch spike peak time (dashed). **E.** Same as D, arranged to reflect the channel layout for Neuropixels probe. Scale = 100 μV. Corresponding Channels in C-E are indicated by colour.

**Figure 4.**
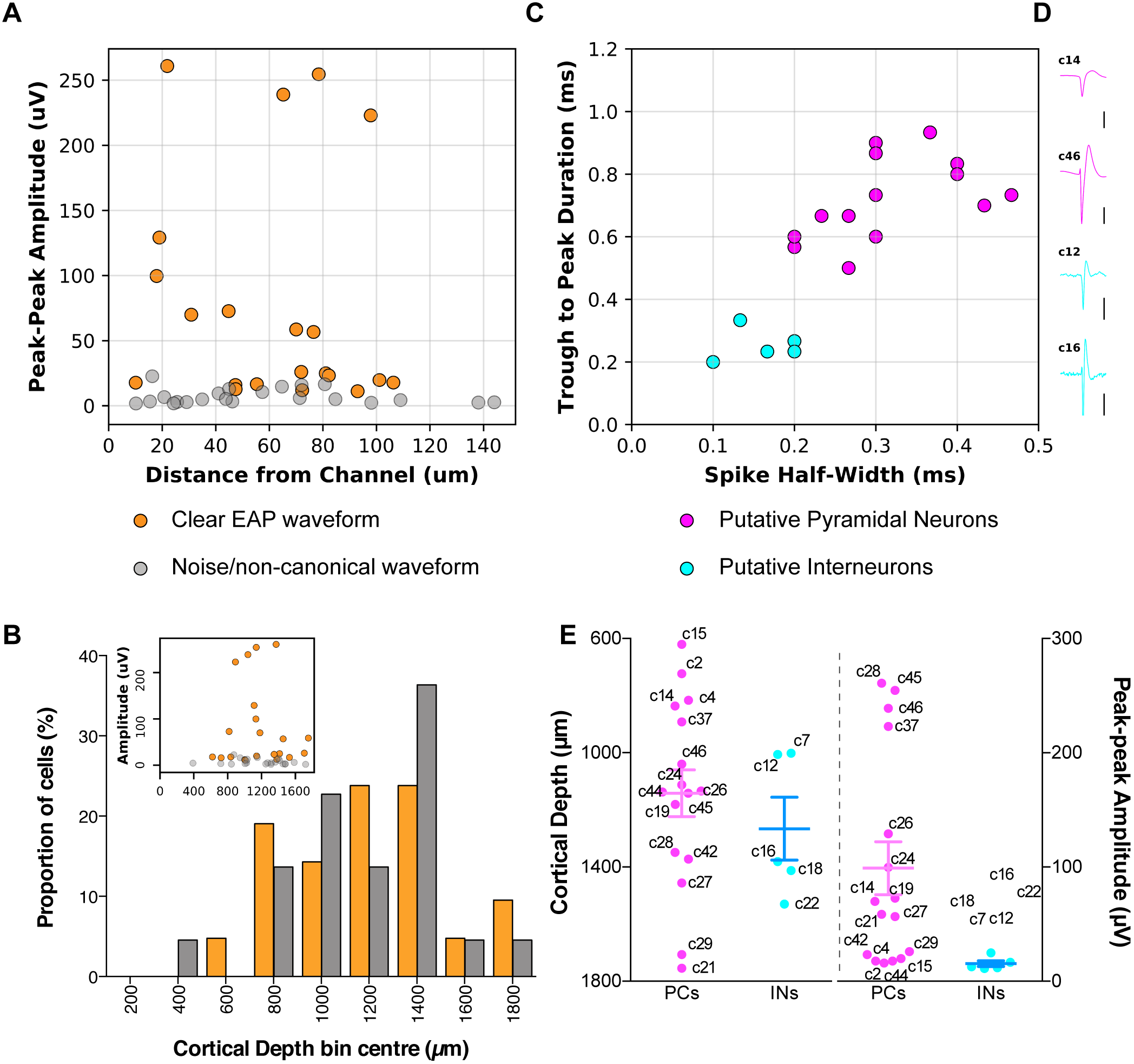
Dataset Summary. **A.** Peak-peak amplitude of PSTA as a function of distance between cell soma and the channel with highest amplitude for every cell recorded. N = 43 cells (21 clear EAP [orange], 22 noise/non-canonical EAP [grey]). **B.** Cortical depth of recorded cells. Inset shows variation in peak-peak PSTA amplitude as a function of cortical depth. **C.** Units with detectable EAP waveforms (orange in 4A-B) could be classified as putative interneurons (cyan) or pyramidal cells (magenta) according to half-width and peak-to-peak duration of the PSTA extracellular spike waveform. **D.** Average spike waveform examples for interneurons (c12, c16) and pyramidal cells (c14, c46). Time window is 2 ms; vertical scale bar is 5μV for putative interneurons and 50 μV for pyramidal cells. Example cells were recorded at depths, respectively, of 1,005, 1,529, 891 and 816 μm, and distances to probe of 93, 55, 97 and 45 μm. **E.** Cortical depth, putative cell type and PSTA peak-peak amplitude for every cell with a clear EAP waveform.

### A summary of the dataset

#### Where did we record?

We recorded a total of 43 neurons with distances of 10-144μm between the patch pipette tip and the closest channel on the Neuropixels probe (Figure 4A; Mean ± SD: 59.6 ± 34 μm). In terms of cortical areas, due to constraints on travel range posed by working with two opposing manipulators, we restricted probe insertion sites along the mediolateral (ML) axis to a 1,261 μm-wide region (minimum to maximum, all coordinates relative to bregma: −3,361 to −2,100 μm). Within this restricted ML axis, we varied anterior-posterior insertion site substantially (−3,526 to 2,527 μm). On the dorso-ventral axis, we inserted the probe deep enough to cover all cortical layers in rat (2,500 to 3,800 μm; cortical thickness in recorded areas ranged from 1,800 to 2500 μm, Paxinos & Watson, 1998). These 3 intervals of stereotaxic coordinates define a stripe of cortex spanning, from anterior to posterior, the following areas: primary motor, primary somatosensory forelimb, primary somatosensory hindlimb and primary somatosensory trunk. Regarding cortical depth, the vast majority of paired recordings were obtained from neurons located at 700 to 1,500 μm from the pial surface (36/43 neurons), suggesting we mostly recorded from neurons in cortical layer (L)5^33^.

We divided our initial sample of 43 neurons into two groups: 21 neurons showed a clear EAP waveform after PSTA, whereas the remaining 22 did not, revealing waveforms that were either non-canonical, or had peak-peak amplitudes below the noise threshold (Figure 4A; see Supplement to Figure 4-1A and *Inclusion Criteria* in Materials and Methods). The laminar distribution of neurons in the two groups was similar (Figure 4B) and bore no discernible relation to distance to the probe (Supplement to Figure 4-3A). Furthermore, the presence of neurons without a clear extracellular signature was not explained by distinct firing rates (Supplement to Figure 4-3C). We will examine possible explanations for putative “dark neurons” in the Discussion section. Until then, we will focus our study on the 21 cells showing clear EAP waveforms after PSTA (orange in Figure 4A-B).

#### What did we record?

Cell types in the brain can be identified by features such as gene expression, morphology, connectivity pattern or properties of the action potential waveform. In our study we had access only to the latter; though not as powerful for parsing out cell types as intracellular recordings, extracellular spike waveforms can at least be used to make a basic distinction between putative inhibitory interneurons and pyramidal cells^34^. To this end, we computed trough to peak duration and negative peak half-width for the spike waveform revealed by PSTA of the extracellular channel with highest peak-peak. This procedure revealed the presence of 5 putative interneurons in our dataset (c7, c12, c16, c18 and c22), which clustered separately from putative pyramidal cells by simultaneously showing the shortest trough to peak and spike half-width durations (Figure 4C, cyan). In our relatively small sample, this proportion (23.8%, n = 5/21) is above the documented ratio of interneurons in rat somatosensory cortex (11.6-12.1%, of all neurons, Meyer et al., 2011). Besides faster kinetics (Figure 4C), EAPs in the 5 putative interneurons in our sample were characterized by lower peak-peak amplitude (Figure 4E; Median, inter-quartile range: pyramids - 64.4 μV, 179.3μV; interneurons - 12.7 μV, 9.7 μV). This difference between pyramids and interneurons was not explained by distance to the probe, as this was not significantly different between them (Median, inter-quartile range: pyramids - 67.6 μm, 57.2 μm; interneurons - 72.2 μm, 35.6 μm; Mann-Whitney U = 31, *p =* 0.495; Supplement to Figure 4-3B). Though our small sample size for interneurons precludes strong conclusions, it is possible that their distinct morphological and biophysical properties contribute to the decreased amplitude of their EAPs, compared to pyramids.

### Detecting patch-clamp spikes on the extracellular probe

The process of analyzing extracellular recording data begins with the detection of action potentials. For paired recordings to be useful in benchmarking or improving spike-sorting algorithms, action potentials recorded in patch-clamp must be detectable on the extracellular recording. To verify this, we detected spikes in the 30 channels closest to each of the 21 neurons as negative crossings of a threshold defined as 7 times the median absolute deviation (MAD) for that channel. To compare the extracellularly-detected spike times with action potentials recorded in patch-clamp, we generated peri-event time histograms (PETHs), computing the times of spikes found in the extracellular channels relative to each patch-clamp spike, for a window of 50 ms around the patch-clamp spike (Figure 5; Supplement to Figure 5C). For some paired recordings, these PETHs showed a high proportion of spike co-occurrence in the patch-clamp and extracellular channels at approximately 0 ms time delay, indicating that the patch-clamped neuron’s spike is being found by the detection algorithm on the extracellular recording (for example, Figure 5A-B, c21 and c14; see others in Figure 5D). For other pairs (Figure 5C, c44; see others in Figure 5D), spikes were less reliably detected on the probe, showing only short peaks (for example, Figure 5D, c27) or barely any peaks at all on the PETH (Figure 5D, c22) around 0 ms. The reason for this is investigated in Figure 5E; the y axis depicts the ratio of extracellular to patch spike detections at the time of the patch spike for the channel closest to the probe. For a probe channel to be judged to be detecting all patch-clamp spikes, it is a necessary condition that this ratio equals or exceeds^2^ 1. The x axis in Figure 5E indicates the amplitude of each cell’s EAP peak relative to the detection threshold (in log-2 fold-change). If a cell’s average EAP peak falls below threshold (left of vertical line), its spikes will always fail to be detected (unless they “ride” occasionally on spikes from background units). The extracellular spike detection ratio for the cells with the highest negative peaks (c28, c37, c45 and c46) suggests these cells had all of their spikes detected. In three cases (c28, c45 and c46), more spikes were detected on the extracellular recording than had actually been fired by the pair-recorded neuron; these false positives likely reflect coincident background activity, which could potentially be separated by spike sorting. To conclude, a total of 9 pair-recorded neurons had >50% of their spikes detected by a simple threshold-crossing algorithm ran on the extracellular trace of the closest channel of the extracellular probe.

**Figure 5.**
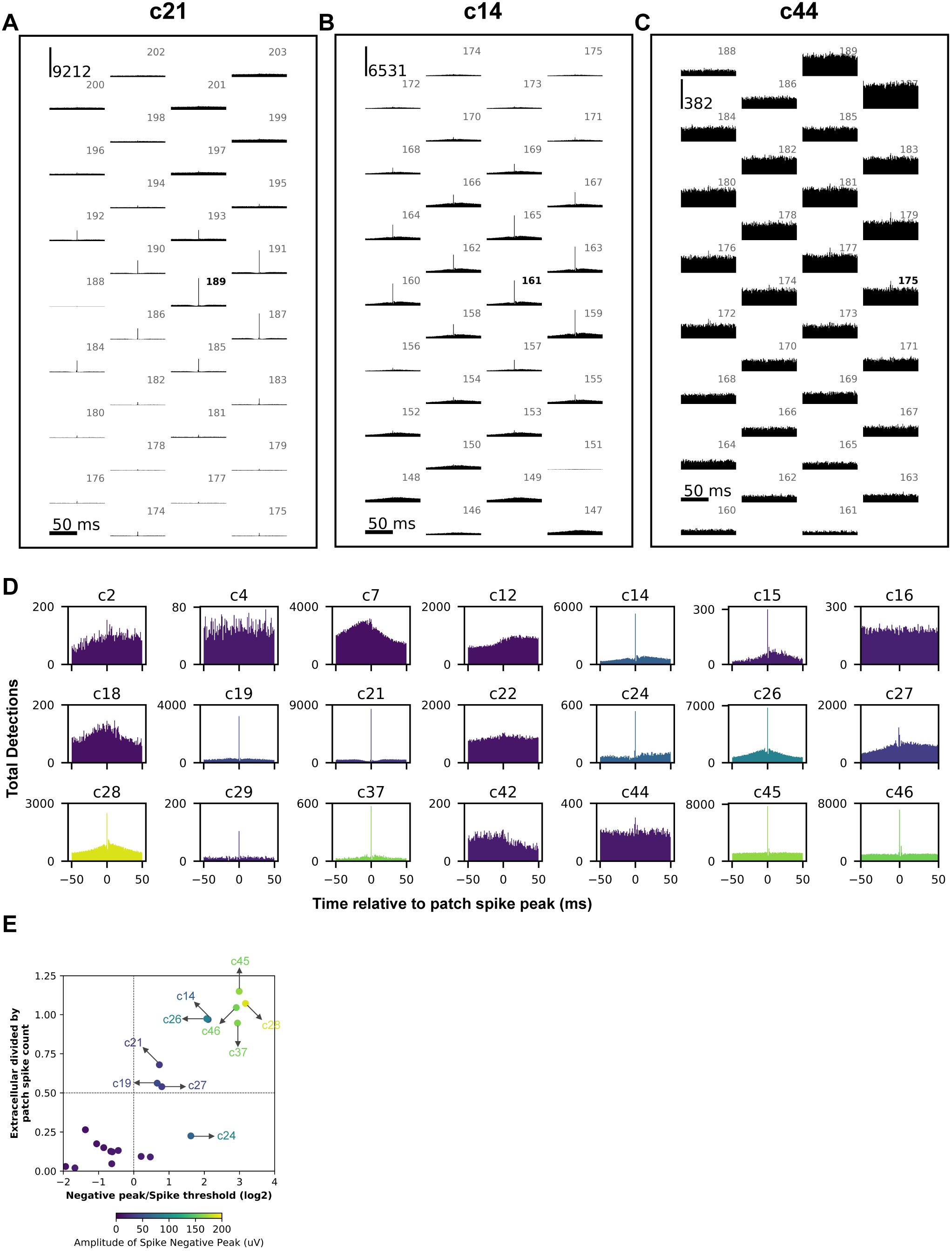
Extracellular detection of the patched neuron’s action potentials. **A-C.** Peri-event time histograms of EAPs detected relative to patch spike times in 1 ms bins centered at 0 ms, for the 30 channels closest to the cell soma. Vertical bar indicates spike count. Channel number indicated above each PETH, closest channel in bold. **D.** As in A-C, for the channel closest to the cell soma. Cells are colour-coded by the amplitude of spike negative peak (colour bar in Fig. 4E). **E.** Y axis: ratio of extracellular to patch spike detections at time of patch spike for channel closest to the probe. X axis: Log2 fold-change between spike negative peak amplitude and detection threshold. Horizontal line indicates detection of 50% of patch spikes; vertical line indicates spike peak amplitude identical to detection threshold.

### Variability in spike features within the same unit

Spike sorting algorithms use extracellular waveform features to cluster spikes and assign them to neurons. Spikes with very different features will in principle have been fired by distinct neurons. But how variable are the extracellular waveform features of spikes fired by *the same unit*? Paired recordings offer a unique opportunity to answer this question empirically with a view to providing constraints for analysis and interpretation of extracellular recordings^18,23^.

To provide an estimate, we investigated the 9 cells in our dataset that showed best extracellular detectability^3^ of patch-clamp spikes. To this end, we looked in the extracellular probe recordings at the 2-ms window either side of each patch-clamp spike time and ran a custom algorithm for extracting spike features (see *Materials and Methods – Analysis*). Despite knowing the precise times for action potential peaks in dual-recorded cells, occasionally other neurons fired action potentials that coincided in time and whose voltage summed with our target neuron’s, distorting spike feature measurements. To skirt this issue, first we extracted features^4^ for the PSTA waveform of each cell. Then, for each pair-recorded spike, we computed the absolute z-score for each feature of that spike, relative to that of the average waveform. By averaging absolute z scores for 8 spike features, we obtained an indicator of how similar/dissimilar a particular spike was from the “pure” spike waveform. Since our goal was to obtain an estimate of within-unit feature variability, we set a strict criterion for spike selection of having an average z score under 1.0. This resulted in an average of 85.4% of spikes being retained for further analysis. All spike waveforms analyzed are plotted in Figure 6A; all those rejected are shown in Supplement to Figure 6-1A.

**Figure 6.**
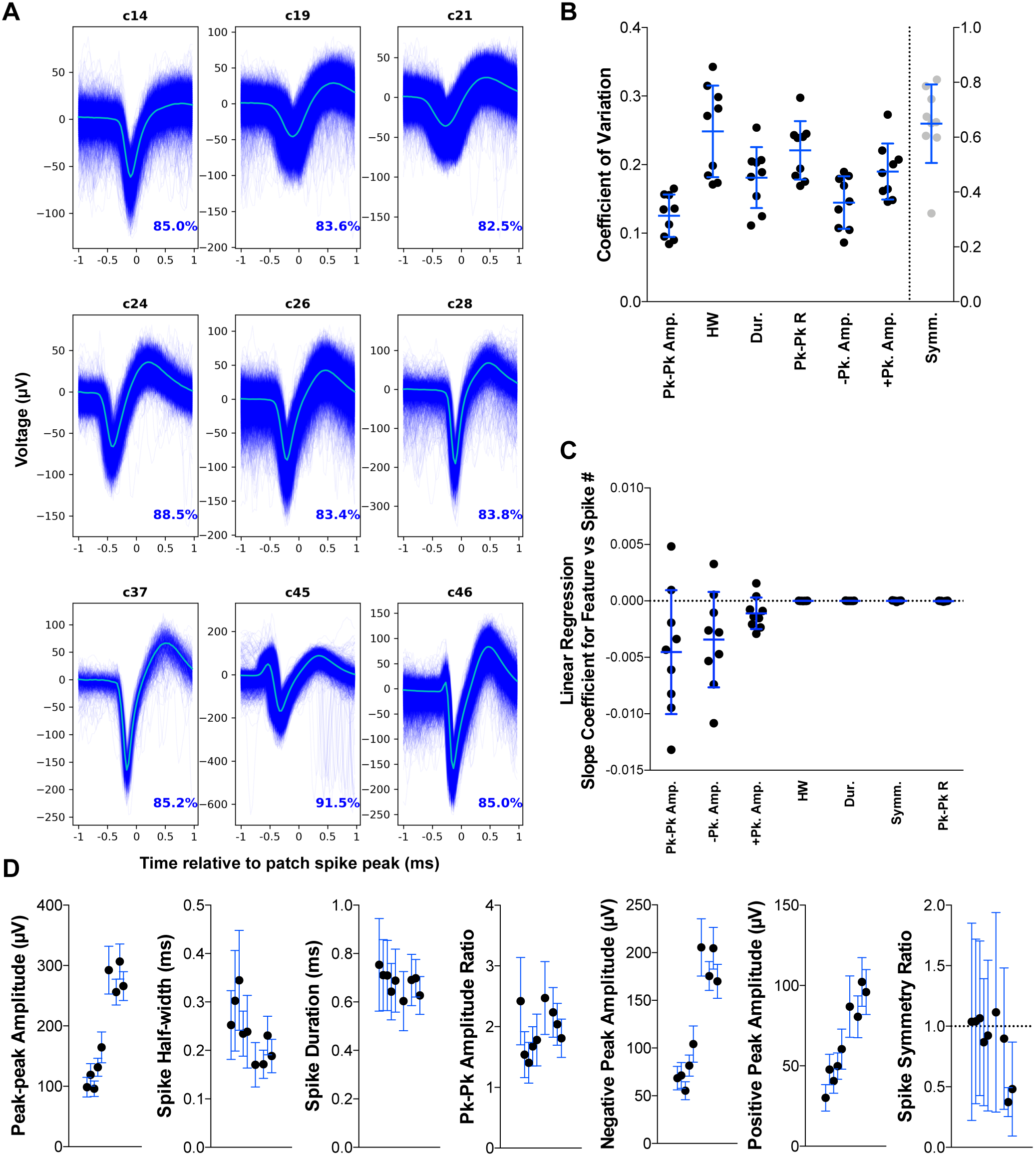
Within-unit variability in spike features. **A.** Extracellular spike waveforms selected for analysis from 9 pair-recorded cells. Average waveform in cyan. Below, percentage spikes analysed out of each cell’s total spikes fired. **B.** Coefficient of variation for 7 spike features in each of 9 cells analysed. Bars indicate grand mean ± SD. Spike symmetry ratio is plotted on right Y axis. **C.** Slope of linear regression between spike number in trial and each of 7 features for 9 cells analysed. **D.** Mean ± SD for spike features in 9 cells.

After plotting each spike feature over time (Supplement to Figure 6-2 to 6-6), it became clear to us that at least for some features, changes occurred at a slow rate over the course of a whole recording. For instance, peak-peak amplitude in c14 (top left, Supplement to Figure 6-2) decreased from approximately 120 μV at the beginning to around 90 μV by the end of the recording. Changes such as this could be explained by a variety of factors, including deterioration of tissue surrounding the probe, drift, or membrane stress and increased “leakiness” caused by the patch electrode. Whilst the first and second factors are part and parcel of any extracellular recording, the latter isn’t and we cannot exclude its contribution. For this reason we attempted to control for slow changes in our analysis of within-unit variability by computing for each feature the average of the *running* mean and *running* standard deviation, instead of averaging each feature outright. From the average and standard deviation we computed the coefficient of variation (CV) for each feature, in every cell, along with the grand mean and standard deviation of all cells. We subsequently report the grand mean ± standard deviation of the CV for every feature analyzed (Figure 6B). In our dataset, the two most reliable features related to spike amplitude: peak-peak amplitude (CV = 12.5 ± 3.1 %) and negative peak amplitude (CV = 14.5 ± 3.8 %). These were followed closely by spike duration (CV = 18.1 ± 4.5 %) and positive peak amplitude (CV = 19.0 ± 4.1 %), then peak-peak amplitude ratio (CV = 22.1 ± 4.3 %) and spike half-width (24.8 ± 6.7 %). Spike duration symmetry was by far the most variable feature (CV = 64.9 ± 14.3 %). Absolute values of mean ± SD for each feature in every cell are plotted in Figure 6D.

Lastly, re-examining the rolling mean and standard deviation plots (Supplements to Figure 6-2 to 6-6), we wondered if any of the spike features we examined changed in a reliable way over time, in this sample of 9 cells. In other words, was there a consistent slope to the change in a particular feature over time? We investigated this by computing univariate linear regression for each feature from the spike index, plotting the slope coefficient in Figure 6C. Of the 7 features analyzed, the 3 that are amplitude-related appeared to decrease slightly over the course of a recording for the majority of cells. For the remaining 4 features, no reliable change was apparent.

### Spatiotemporal dynamics of the extracellular action potential waveform

Spikes from a single neuron can be detected by multiple channels on extracellular probes, especially in probes with high channel density (Figures 2-4). This feature is exploited by spike sorting algorithms^14,15,17,27^, and therefore knowing more about how voltage spreads over space and time during a spike may contribute to sorting units more accurately. Moreover, spatiotemporal structure in spike waveforms across many channels may reveal other potentially interesting sources of information to exploit, such as the contribution of different subcellular components to the extracellular signal, or the location and direction of propagation of action potentials.

The leftmost panel of Figures 7A and B is an attempt to depict the rich spatiotemporal dynamics of action potential spread. We calculated the first derivative of voltage over time (normalized to its absolute maximum across the probe) for the PSTA of each channel in a single column of the probe (192 channels, 40 μm spacing row to row), color-coded and plotted it over time. Red values indicate a decrease in voltage signal, blue an increase and white no change. For every channel in the column, we can therefore track the progression of voltage over time, for a spike. The right-hand panel in Figures 7A and B shows the waveform of each spike within the dashed region indicated to the left. In c24, we can see that the earliest change in voltage for channel 218 (indicated on left; corresponding magenta trace at right) is a negative shift. After reaching the negative peak (white pixel), this is followed by an increase in voltage (blue), corresponding to the repolarization phase of the spike. After reaching a positive peak (white), voltage decreases again to near baseline levels (red). Channels near the soma follow a similar progression but delayed in time. As we move upward along rows, a different pattern is observed (both can be seen in c24 and c21), where the earliest phase is a positive voltage deflection, followed by a negative peak and a second positive deflection, completing a “triphasic” spike. As one journeys farther up along the probe, the second positive peak disappears. Spikes with triphasic or positive-then-negative patterns have previously been implicated with the apical subcompartments of the dendritic tree^25^; it is interesting to note that for c21 the early positive deflection can be seen at rows over 480μm away from the probe. It is possible that for this particular recording, the apical dendrite was in close parallelism to the axis of insertion of the probe. A final point worth noting in c21 is that the earliest negative peak is *not* at the channel closest to the soma, but rather a couple of rows below it. One possibility is that this “early” channel was detecting the electrical field produced by the axon initial segment. Similar plots for all remaining cells are presented in Supplements to Figure 7-1 to 7-4.

**Figure 7.**
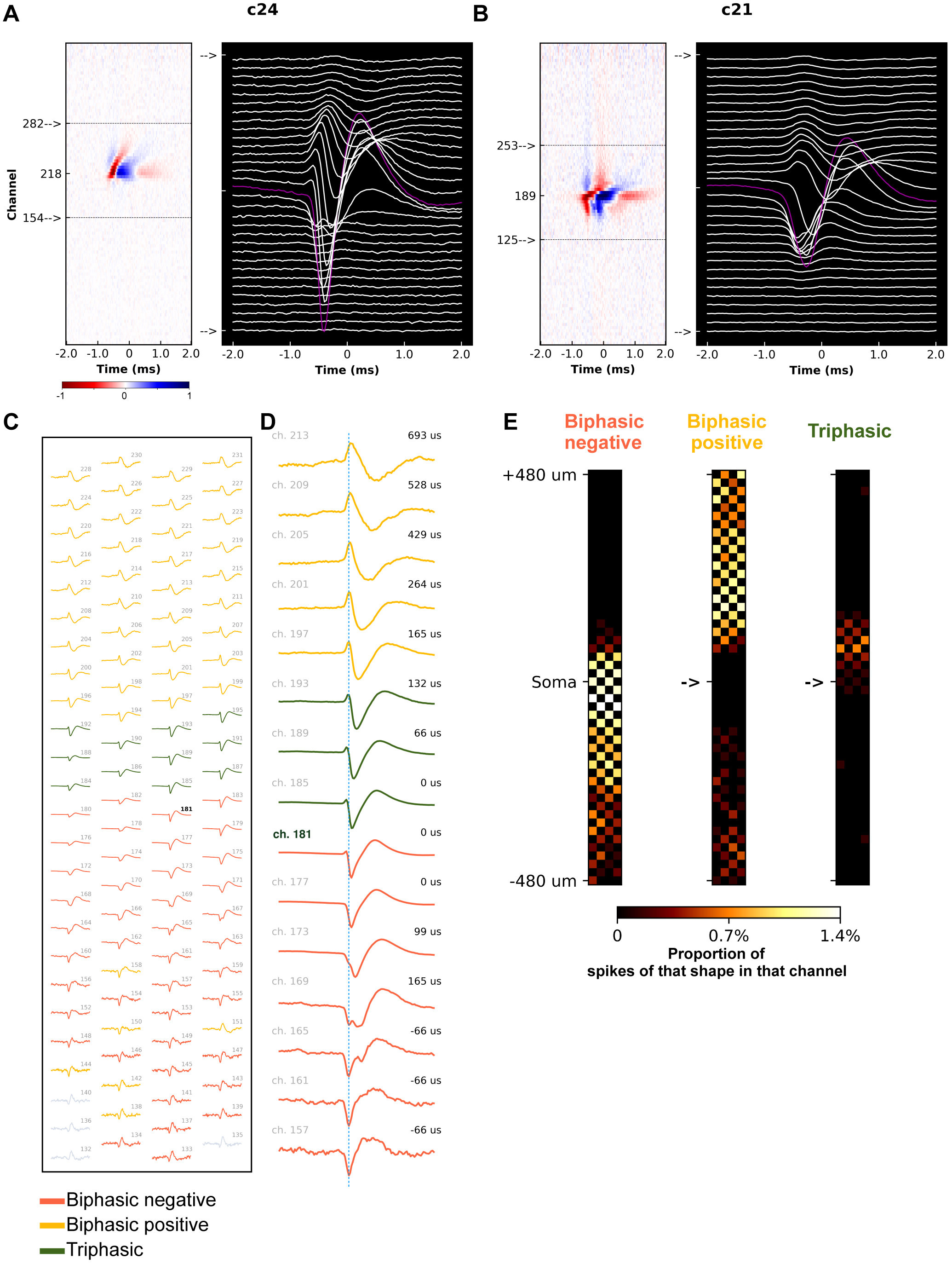
Spatiotemporal dynamics of the extracellular action potential waveform. **A, B.** Left: Heatmap of normalised derivative of voltage over time of PSTA for each channel in a single probe column. X axis is time relative to patch spike peak. Colormap is negative (red) to positive (blue). Right: PSTA waveforms for 32 channels delimited by dashed box (left). Channel closest to the soma in magenta. C. PSTA waveforms for 100 channels on c46. Waveforms were classified as: biphasic positive (yellow), triphasic (green) or biphasic negative (red). Grey is unclassified. Voltage scaled row-by-row for visibility. Channel number indicated at top right of each channel (bold for closest). **D.** PSTA waveforms for 15 channels on the same probe column as the soma of c46. Traces are aligned by patch spike peak and dotted line is the earliest negative peak detected for c46 PSTAs. **E.** Distribution of waveforms of each shape over 100 channels either side (DV) of the recorded cell, for 10 putative pyramidal neurons (708 PSTA waveforms).

Based on the variety of waveforms observed in Figure 7 and its supplements, we asked if, in line with previous work^25^, there was any regularity in terms of where, relative to the cell soma, these patterns were typically observed. To this end, we implemented a very simple classification of PSTA waveforms into the categories of *Biphasic negative* (negative peak followed by positive peak), *Biphasic positive* (positive peak followed by negative) and *Triphasic* (short positive peak, followed by negative, ending with a more prominent positive peak), based on order of crossings of positive and negative thresholds by the first derivative of the voltage over time. Figure 7C shows the PSTA waveforms for 100 channels (centered on the closest channel to the soma, in bold) for c46. Waveforms are color-coded according to their classification, and the voltage axis is scaled per row, to improve visibility of the signal. As for all our recordings, channel numbers follow a progression from deep in the brain (low numbers) to its surface (high numbers). As can be seen from Figure 7C, biphasic negative waveforms (red) tend to appear near to and below the soma, whereas biphasic positive spikes are seen towards the cortical surface (yellow). In between, there are triphasic (green) waveforms. Figure 7D shows example traces of each waveform class across a single column of the probe for c46. We plotted such maps for every one of the 10 putative pyramids with PSTA amplitudes > 50 μV (Supplement to Figure 7-5/6). We next classified PSTA waveforms and quantified their distribution over 50 channels either side of the putative site of the soma. A total of 1,000 PSTA waveforms were analyzed, corresponding to 100 waveforms from each of 10 pyramidal cells. Out of all cells, in 200 waveforms, no spike could be detected. Out of the remaining 800, 708 spikes could be classified into one of three types. Biphasic negative spikes accounted for an average of 46.3 ± 11.9% of classified PSTA waveforms (Figure 7E, Supplement to Figure 7-5, 7-6 and 7-7, 7-8A) and were mainly observed at the soma and locations of 160 μm below to 80 μm above it (dorso-ventral axis). Biphasic positives compounded 46.9 ± 10.0% of waveforms per cell and were found mostly above the soma, extending up to 480μm towards the surface, whereas Triphasic waveforms were the rarest (6.9 ± 6.4% of spikes per cell) and restricted themselves to a short segment tiling the space ~100 μm above the soma.

We stress that our classification algorithm was highly simplistic, missing out on finer waveform distinctions (e.g. Figure 7D ch165 vs ch177). However, bearing in mind the orientation of our probe insertions, even this crude approach captures a tentative regularity which is consistent with previous work: biphasic negative spikes are mainly found near the soma and proximal basal dendrites; biphasic positives can usually be observed at distal segments of the apical dendrite and lastly, triphasic waveforms (positive-negative-positive) tend to locate to the proximal apical trunk^25^.

## Caveats and limitations

### Distance estimates

Although we calibrated our approach to measure distance between probe and patch pipette tip extensively, by necessity all such tests and calibrations were conducted outside the brain. We therefore assumed that the rigidity of the probe and patch pipette would be sufficient that their penetrating movement would not be disrupted significantly by brain tissue and therefore affect our measurement of distance. Furthermore, since we did not recover histological data (probe tracks and neurobiotin fills), any bending of the probe due to encountering a volume of tissue with higher rigidity (for example a large blood vessel) would have been invisible to us during experiments. In our experiments, the only sign of this occurring would have been recording a series of neurons with lower PSTA peak-peak amplitudes than expected by our estimated distance to the probe (for example estimating that we were 30 μm away from the probe when we were in fact 100). Examining Figure 3A we can see that a subset of 12 neurons (grey points) located at less than 60 μm from the probe did not show clear EAP waveforms. Although factors such as cell orientation relative to the probe and morphology can also greatly affect extracellular peak-peak-amplitude, we cannot exclude that probe displacement within the brain affected some of our distance estimates. However, our findings are consistent with the broad consensus that 60-80 μm is the maximum radius from which extracellular probes are able to record action potentials (Figure 4A; refs. 18, 23). Furthermore, for 70% of recordings with clear EAP waveforms, our estimate of the channel closest to the soma was off by 5 channels or less, suggesting our distance estimates were reasonably accurate. We stress that the goal of our technical approach to paired recordings was to develop a method that would enable us to greatly increase the throughput of these challenging experiments, returning a reasonably accurate *estimate*, rather than exact measure, of distance from probe channel to soma.

### Intracellular solution

A second point we would like to draw attention to is the fact that we opted to use high-potassium current-clamp intracellular solution for our recordings. There were two reasons driving this decision; first, we wished to be able to break in and record in whole-cell configuration, in situations where we successfully obtained a giga-seal; second, dual recordings from the same neuron *in vivo* are time-consuming, labour-intensive and technically-challenging. The degree of experimental difficulty is further compounded by the fact that to be useful, these recordings must contain a reasonable number of spikes. In whole-cell configuration, this is never an issue since the experimenter can inject current into the cell and drive it to fire action potentials. However, in cell-attached mode it may not always be possible to drive a “silent” neuron to fire. We reasoned that in such situations, high-potassium intracellular solution would help depolarize a neuron and increase the likelihood of recording spikes. Whilst this did indeed prove to be a productive approach, it means that firing rates in channels close to a dual-recorded neuron’s soma will be artificially inflated by potassium-rich medium. We therefore advise collaborators to take this point into account when planning analysis and exploration of our dataset.

## Discussion

### Summary

We built a software-assisted dual recording rig^23^ (Materials and Methods) that allowed us to patch-clamp neurons at distances close to a Neuropixels dense CMOS probe^10^. This enabled us to record a total of 43 neurons, from which 21 showed clear EAP waveforms on at least one channel of the probe, producing a publicly-available “ground-truth” dataset that can be used for development and validation of improved spike sorting algorithms. In accordance with previous reports^18,22,23^ we find that distance is the major factor determining the peak-peak extracellular signal amplitude, though cell type-specific factors may also play a role. Our approach enabled us to separate with high precision the activity of dual-recorded units from cells spiking in the vicinity. We were thus able to provide an estimate of within-unit variability for a few commonly-used spike features. Finally, we highlight the richness of spatiotemporal signatures present in extracellular recordings, which is emphasized in high-density probes^10,11^.

### A publicly-available “ground-truth” dataset

The full dataset (http://bit.ly/paired_recs),metadata, accompanying information and analysis code is available online (http://bit.ly/paired_git); the interested reader can obtain any further information by contacting the corresponding authors. Validation datasets (present study; refs 18 and 23) are invaluable for benchmarking and improving analysis algorithms and provide empirical answers to questions about neuro-technological development. Moreover, tethering a complex signal (extracellular voltages over time in hundreds of channels distributed in space) to a simpler one (patch-clamp recordings from a single neuron identified in 3D space) helps us better delineate signal from noise – and potentially uncover novel sources of information, previously treated as noise.

### “Dark neurons”

There is a long-standing debate over whether we are missing neurons with extracellular recording. This stems from a discrepancy in the number of neurons reported to be firing action potentials when using optical (~50%) compared to electrical (< 10%) recording techniques^35,36^. However, any estimate on the number of neurons an electrode should detect depends on knowing how far away a neuron’s spike can be detected. The distance limit for recording EAPs has been estimated by ground truth^18,22,23^ and modeling studies^36–39^ to be around 50 μm, though reports exist of recording unexpectedly large (>50 μV) EAP waveforms at greater distances from the probe^18,40^ (present study, see 3 cells in Figure 3A with EAPs > 200 μV and distance 60-100 μm). Conversely, we found 14 neurons within the estimated range of extracellular detection (50 μm) that produced small or undetectable EAPs even after averaging thousands of spikes (Figure 4A). It has been suggested that factors beyond distance can also contribute to EAP detectability: for instance, bias of extracellular recordings to highly-active neurons and some cell types producing weaker and more spatially-localized extracellular signatures^35,41^. We can rule out the former in our present study, as we separately monitored spiking activity with patch-clamp, finding no correlation between firing rate and EAP amplitude for extracellularly-detected and -undetected neurons (Spearman r = 0.115, p = 0.464; Supplement to Figure 4-3D). As for the latter, the small sample of interneurons in our study tended to produce EAPs of smaller amplitude than pyramidal neurons at equivalent distances (Supplement to Figure 4-3B). Beyond these small pieces of information, it will not be our study that ends the debate. Due to methodological limitations (see *Caveats and Limitations*), we cannot safely call the subset of 14 neurons lacking clear EAP waveforms at < 50 μm from the probe “dark neurons”. All we can provide at this point is further information about the sessions in which each neuron was recorded (Supplement to Figure 4-3C). Out of a total of 15 recording sessions, 6 contained a mix of neurons with and others without clear extracellular spike waveforms after patch spike-triggered averaging (light blue), and 3 resulted only in neurons with clear EAPs (dark blue). Two sessions initially produced neurons that could be detected extracellularly, but ended with cells that could not (light red), whereas the remaining 4 sessions produced only undetected neurons (dark red). The latter two types of session are consistent with potential errors of manipulator alignment or bending of the probe after entering the brain (see *Caveats and limitations*), which could explain the lack of clear EAP waveforms in the resulting neurons. Therefore, to obtain a more accurate list of potential “dark neurons”, we should exclude 1) cells recorded in red and light red sessions and 2) cells from which we recorded less than 200 spikes, as this number might not have been high enough to isolate a cell’s firing from background activity. Narrowing down our list by these two criteria results in 7 cells: c2, c3, c13, c15, c23, c25 and c36. We advise any investigators wishing to mine our dataset for “dark neurons” to confine their explorations to these recordings.

### Reliability of EAP waveform features

The dual-recording setup allows experimenters to identify the spikes fired by a single unit. In a way it is - by perhaps not so great a margin - the most labor-intensive and time-consuming method for manual spike sorting, but also the most accurate. We exploited this advantage to address the issue of variability in features of spikes fired by the same unit. From the features we examined, our results suggest that peak-to-peak amplitude, negative and positive peak amplitude and peak-peak duration are the most reliable qualities of an action potential, with standard deviations under 20% of the value of their mean. Spike half-width and peak-peak amplitude ratio were somewhat more variable, but their standard deviations were still under 30% of the value of their mean. We hope these estimates are useful but we highlight that they were obtained in urethane-anaesthetised rats and cannot rule out that variability is different in awake recordings, or that our simultaneous patch-clamp recordings affected the variability of certain features.

### Spatiotemporal properties of the EAP waveform

One of the most exciting features of the new generation of CMOS probes is their high channel count and density, enabled by multiplexing and advances in microfabrication^12^. The short spacing between channels (20 μm) is on par with the spatial scale of changes in neuronal morphology features. Given that a neuron’s subcellular compartments produce distinct extracellular spike signatures^25^, we wondered if this could be exploited to extract gross morphological information (such as orientation, or the type of compartment nearest a channel) from extracellular recordings. Our highly-simplistic approach produced results consistent with models^25^ based on previous ground-truth experiments^18^. We suggest it would be of great interest to acquire a further ground-truth dataset with CMOS probes where the morphology of a dual-recorded neuron is recovered. These experiments are challenging and likely to require high numbers of cells, but the potential utility of estimating gross morphology - or the position in 3D space of a unit’s soma – would be immense.

## Potential projects for collaboration

We aimed for this manuscript to perform the function of a “data descriptor”, explaining the methods and restricting ourselves to mainly descriptive analysis to familiarize the reader with our dataset. In releasing the dataset publicly, our hope is that it will help neuroscientists currently developing spike sorting and analysis algorithms, but also that it will be used to explore further questions. In particular, we hope to foster peer-to-peer synergy and open collaboration through our sc.io repository (http://bit.ly/paired_git), with a view to authoring high-quality, reproducible follow-up publications jointly with a community of interested scientists.

Here we outline and explain potential projects for collaboration. These projects are set up as individual branches of the repository which anyone can contribute to or fork to start their own independent work. Interested scientists are also free - and in fact, encouraged - to take the lead and pose new questions by proposing branches for the sc.io repository.

### 1. Studying synaptic connectivity with dense CMOS probes

One of the most exciting features of high channel-count/density probes is that the increase in channel number produces a dramatic increase in likelihood of finding connected pairs^42^. This will go a long way to advance our ability to study synaptic modulation, dynamics and plasticity in behaving animals under different brain or behavioral states. By isolating units with patch-clamp, our dataset offers a solid testing ground for examining cell-to-cell interactions. There are 5 whole-cell recordings in it where one could potentially detect sub-threshold post-synaptic potentials and search for putative pre-synaptic units, mapping microcircuits across an entire cortical column. A second feature of interest is that we obtained a mix of dual-recorded putative interneurons and pyramidal cells, enabling examination of excitatory and inhibitory interactions between cell types.

### 2. Exploring axon terminal and field post-synaptic potentials (AxTPs and fPSPs)

Axon terminals generate low-amplitude electrical fields that fall far below the noise threshold. However, by averaging hundreds or thousands of spikes, if a unit’s axon terminal happens to fall close enough to an extracellular channel, AxTPs can be revealed^43,44^. A potential confound is to ensure these small signals are active axonal potentials and not passive detection of the somatic action potential’s electrical field, attenuated by distance. The distances covered by the shank of the probe used (3.82 mm) ensure that plenty of channels will be far enough from a unit’s somatic location that AxTPs may be revealed. fPSPs may also be investigated at these distal locations, by studying the LFP band. The high density and channel count of CMOS probes increases the likelihood that axon terminals or distal dendrites will fall within the detection range of a channel, making highly-localized signals such as AxTPs and fPSPs more tractable. Ultimately, these signals could offer strong mechanistic support for inferences about unit-unit connectivity.

### 3. Extracting orientation and gross morphological features from extracellular recordings

In a landmark ground-truth study, Henze and colleagues (2000) recorded from the same neuron using tetrodes and intracellular recordings, obtaining morphological reconstructions for a subset of these cells. Follow-up modeling studies intricately linked morphological and conductance features to aspects of the EAP waveform^25,26^. In the present study, we used a crude approach to find regularities in waveform shape that related to estimated neuronal orientation, achieving results in agreement with Gold and colleagues (Figure 7). Given a) known constraints on the attenuation of electrical fields with distance in the brain, b) previous biophysical conductance modeling studies (Gold et al., 2006) and c) the high channel density of CMOS probes, can we begin to approximate gross morphological features, neuronal morphology orientation or location in 3D space for extracellular units? If not, what data are we missing to achieve this? Imaging-based approaches to recording neural activity offer the ability to relate spikes to identified neurons and their cellular and neurochemical properties^45,46^. Unlocking this possibility for extracellular recordings would be an extraordinary development, bridging levels of analysis from molecules to behavior and potentially aiding the sorting process by offering extra clustering dimensions (e.g. cell-type, immediate-early gene or ion channel transcripts).

### 4. Do “dark neurons” share common features?

There are 7 neurons in the present dataset whose EAPs could not be recorded by the extracellular probe, even though they were within the estimated range of detection. For these 7 paired recordings, we have excluded misalignment or probe bending as alternative explaining factors. Are there features common to these neurons (or brain/local network state during their recording sessions) that can be gleaned from the patch-clamp or extracellular recordings?

### 5. Are there cell type-specific differences in EAP amplitudes that affect detectability in extracellular recordings?

Conversely to question 4, are there common features to the neurons in the dataset that showed the highest peak-peak EAP amplitudes? We note that not all neurons with high EAP amplitudes were close to the probe; 3 cells with peak-peak above 200 μV were located 60-100 μm away from the closest channel, “outperforming” several neurons located closer (Figure 4A). Other studies^18^ also reported neurons with higher peak-peak amplitudes than would be expected from their distances. Are there biophysical or morphological properties that make certain types of neuron more detectable in extracellular recordings?

### 6. Benchmarking automated sorting algorithms

From the point of view of reproducibility, we would like spike sorting to become a fully automated process. This is a very challenging goal, but great progress has been made towards it^17,27,47^. It would be useful to compare and contrast performance across algorithms on the same datasets, noting situations where each does well or underperforms with a view to learn, develop and optimize consensual frameworks.

### 7. The “psychophysics” of human spike sorting – what are the features or steps of manual sorting and curation that show the highest and lowest inter-operator variability?

Manual and human curation steps in spike sorting are widely known to show high inter-operator outcome variability^24,48^. We would be interested in running a large-scale study where hundreds of human operators start from the same recording (either a raw recording, or the intermediated output of a semi-automated algorithm run on a ground-truth recording) and post their final results online to a database. It would be interesting to consensually collect additional data on variables such as level of experience on manual sorting, individual notes on which features drove specific decisions to merge or split, and potentially even eye-tracking or “reaction time” (how long did an operator take to make a specific decision and is this representative of its difficulty?) data. The ultimate goal would be, as we put it, to gain understanding about the human psychophysics of the sorting process – are there perceptual, cognitive or human factor biases that enhance or impoverish performance? Can we “protect” against them programmatically? Can we learn from strategies used by experts and implement them in spike-sorting algorithms?

## Epilogue: an experiment in collaboration and publication.

### A bright future for Neuroscience?

Technological developments and the rise of large-scale neuroscience research initiatives ^49–52^ are often said to have triggered an age of “big data” in neuroscience. Regardless of one’s views on the merits of drawing comparisons with particle physics, the fascinating reality is that neuroscientists can nowadays image 10,000 neurons with two-photon microscopy ^53^, record hundreds to a few thousand neurons with extracellular probes at millisecond resolution^10,11^, map synaptic input onto a single identified cell ^54–56^, and use clearing/tracing techniques to reveal sub-micron resolution whole-brain projection maps for single neurons. Some of these technologies can be combined, and they are not even an exhaustive list of the powerful tools available to the 21^st^ century neuroscientist. As a consequence, neuroscience is producing more data than ever before and the tools available to us are - by more interpretations of the term than one - <u>awesome</u>.

But is this all that matters? Are we walking the path of righteousness or strolling down the yellow brick road?

### Curb your enthusiasm

We are as excited as the most optimistic amongst our peers about the tools and data becoming available, but feel strongly compelled to raise a few points about how to handle this opportunity and about the wider scientific context in which it is arising.

First, we strongly believe that current paradigms for publication are obsolete, straining under the weight of gigantic datasets and going short of breath to catch up with the speed at which they’re acquired. We not alone in expressing concern about what to do with and how to interpret the current deluge of neural, genetic and behavioral data, as well as in expressing the benefits that a culture of open science and data sharing could bring to neuroscience ^57–66^. Notably, some of the large-scale initiatives mentioned above are highly collaborative and publish through entirely different models, sharing and promoting the re-use of high-quality datasets worldwide. Many academic neuroscience labs should perhaps take note. Second, it is of the utmost importance that scientific strides are supported by solid ground, such that scientific progress is not replaced by the superficial appearance thereof. Third, if we must measure progress and scientific quality, let us tether such indices to real scientific impact and desirable^5^ behavior^66^ instead of the “h-factor”, “Impact Factor”, re-tweets, downloads and other dubious, self-serving and scientifically-bankrupt “metrics” widely-adopted by hiring committees and the commercial publishing industry. Fourth, scientific experiments are a noble but deeply human activity, as fallible and imperfect as the scientists conducting them; the last line of defense of the scientific method is its powerful self-correcting nature, which relies critically on transparency.

For these reasons we believe it is high time that the neuroscience community explored and demanded tools for publishing, collaboration and sharing that match the modernity of the tools it uses for acquiring data. In particular, the neuroscience publishing pipeline of the 21^st^ century must emphasize data sharing and re-usability, reproducibility in processing and analysis algorithms, complete transparency in procedures and materials and finally, adoption of platforms that can support complex, multi-modal data and interactive visualizations.

### Requirements and Implementations

Our first intent in this Epilogue was to raise awareness for the need to develop tools to improve the current ecosystem for sharing, handling, and reporting neuroscience data. Second, we aimed to set out requirements and specifications for what a modern and improved environment should look like. Focusing on requirements instead of prescriptions recognizes that there are many possible implementations that abide by them, keeping the field open for alternative solutions to emerge prior to optimization and consolidation. Furthermore, redundancy in open science tools is important, as open resources may not always remain so. Our own implementation is simple and based on a combination of easily-accessible data hosting services, a free code repository and an open-access preprint server. We will outline how our particular implementation for this project addresses the concerns and requirements we set out.

1. **Data Sharing and re-usability**. The full raw dataset and supporting metadata are available for download through a freely-accessible hosting service. We have provided in the sc.io repository (http://bit.ly/paired_git) instructions on how to download the full dataset or subsets of recordings. In this manuscript on BiorXiv we explain the methods and provide context and caveats to our data, making clear any factors conditioning re-usability. Further clarifications are possible through email contact with the corresponding authors (AMS and ARK).
2. **Reproducibility and transparency in algorithms, procedures and materials.** All code used to process and analyze data and produce figures is freely available at the sc.io repository, ensuring the reader can run it on the raw data and obtain the same results. We welcome any feedback on the code and analyses, especially mistakes that may have eluded us. GitHub code repositories have been designed exactly with this in mind, providing state-of-the-art version-control. The reader can “raise issues” and submit “pull requests” if they have a proposal to fix or improve any code. As for materials and procedures, since preprints on BiorXiv have no length or formatting requirements, we sought to provide as much procedural and material detail as possible in the Methods section, so that scientists interested in conducting similar experiments can work out how to do so from the information we provided.
3. **Dealing with the size and complexity of datasets and the fast pace at which they’re acquired.** In principle, we have finished data acquisition for this project. However, should we perform new experiments, new data can be uploaded and disseminated quickly and existing/new code ran on it to update figures and results. A new version of the manuscript can be uploaded on BiorXiv, which also addresses history by keeping prior versions available. Queries and feedback can be provided with great responsiveness through the GitHub repository, generating fast cycles of assessment, error-checking and review. As we outlined before, our simple experiment generated a complex dataset that can be explored in many ways. By adopting a principle of “open input” through the sc.io repository, we distribute the challenges posed by our dataset to a talented and interested community in the hope that together as a team we can address the underlying complexity productively.
4. **Quality, impact and credit assignment**. Future contributions from the community – to this manuscript or any follow-ups – are publicly-visible and entirely transparent on the GitHub platform as the history of “commits” and activity on the repository, and as such are easy to build into consensual authorship. Every interaction (raising issues, submitting pull requests, committing code, providing comments and suggestions) is trackable, making credit assignment straightforward since each author’s contribution is self-evident and public. Since all work is open, quality can be assessed through scrutiny from the participating community and by soliciting peer review through preprint journal clubs (such as https://prereview.org/) and other contacts. Impact (as per our operational definition) can be measured by tracking how often the dataset is re-used by other authors in publications. This can be extended to analytical and conceptual contributions from ourselves or our collaborators, i.e., by measuring how often a branch of the repository is “forked” (copied by someone else to use as a starting point on that person’s own analysis/idea).
5. **Dealing with rich data, avoiding “flat” visualizations and supporting interactivity**. GitHub repositories can also be used to host websites showcasing interactive content. One approach to interactivity and rich data visualizations is to host a repository with all the code used for analysis and figure generation, and use the media outputs of that code to build a website that displays it elegant and interactively. This has been pioneered and used to great effect by the York group (see https://andrewgyork.github.io/ for publications and a template). A very desirable future development would be for preprint servers to accept submissions not just in pdf but also html and xml. Jupyter (http://jupyter.org/) and other electronic notebooks are also excellent approaches to provide rich, reproducible and clear visualizations. A very interesting development is Stencila (https://stenci.la/), a software project dedicated to developing an open source office suite for reproducible research. A key advantage of Stencila is lowering the technical entry barrier for scientists not as comfortable with code to be able to produce reproducible interactive documents.
6. **Peer-review**. A criticism commonly heard when discussing publishing on preprint servers is the lack of peer review. By sharing our code, hosting discussion and fostering collaboration on a GitHub repository, we provide access to our analysis routines, enabling peer review as comments, issues and pull-requests. Furthermore, peer review can be solicited through preprint journal clubs and direct requests to recognized experts. With due permission, this feedback can be compiled and posted either on BiorXiv or on GitHub, and potentially even given a DOI (https://prereview.org), making it discoverable and citable. Authors and reviewers can engage in dialogue through each of these platforms and publicly set out plans of action to address criticism and improve manuscripts.

## The experiment begins

We have provided and backed up with arguments our perspective on why neuroscience needs better paradigms for publishing and working openly. We have also proposed brief specifications for this. We finish with an example of an implementation which is deployable right now, relying on easily-accessible tools. For now, we look forwards to testing our implementation, initiating a worldwide distributed collaboration, and getting feedback on our electrophysiology work.

We thank you, the reader, for being patient and getting this far.

This is not an ending, but the beginning of an exciting experiment.

## Footnotes

1 In the sense that the spike times of a verified single neuron (unit) are known.

2 In the case where the channel picks up spikes from a unit that is not the pair-recorded cell.

3 Further inspection of c27 revealed the presence of a highly active “background” neuron, which can partly be inferred by the short “head to shoulders” ratio of its PETH (Figure 5D). We replaced it with c24, which was the next best-ranked neuron in detectability, for cells with negative peaks above threshold.

4 Peak-peak amplitude, half-width, duration, peak-to-peak amplitude ratio, latency to patch spike, duration symmetry (duration of hyperpolarising vs depolarising phases of centred on negative peak), negative peak amplitude and positive peak amplitude.

5 A difficult concept to define, but an interesting operational definition is to answer the question “How much did this work change the way I do experiments or interpret them?” (from discussion at 1^st^ SWC Open Neuroscience Workshop, London UK, 25/05/2018).

