## Supplement to Figure 4 for "Recording from the same neuron with high-density CMOS probes and patch-clamp: a ground-truth dataset and an experiment in collaboration"

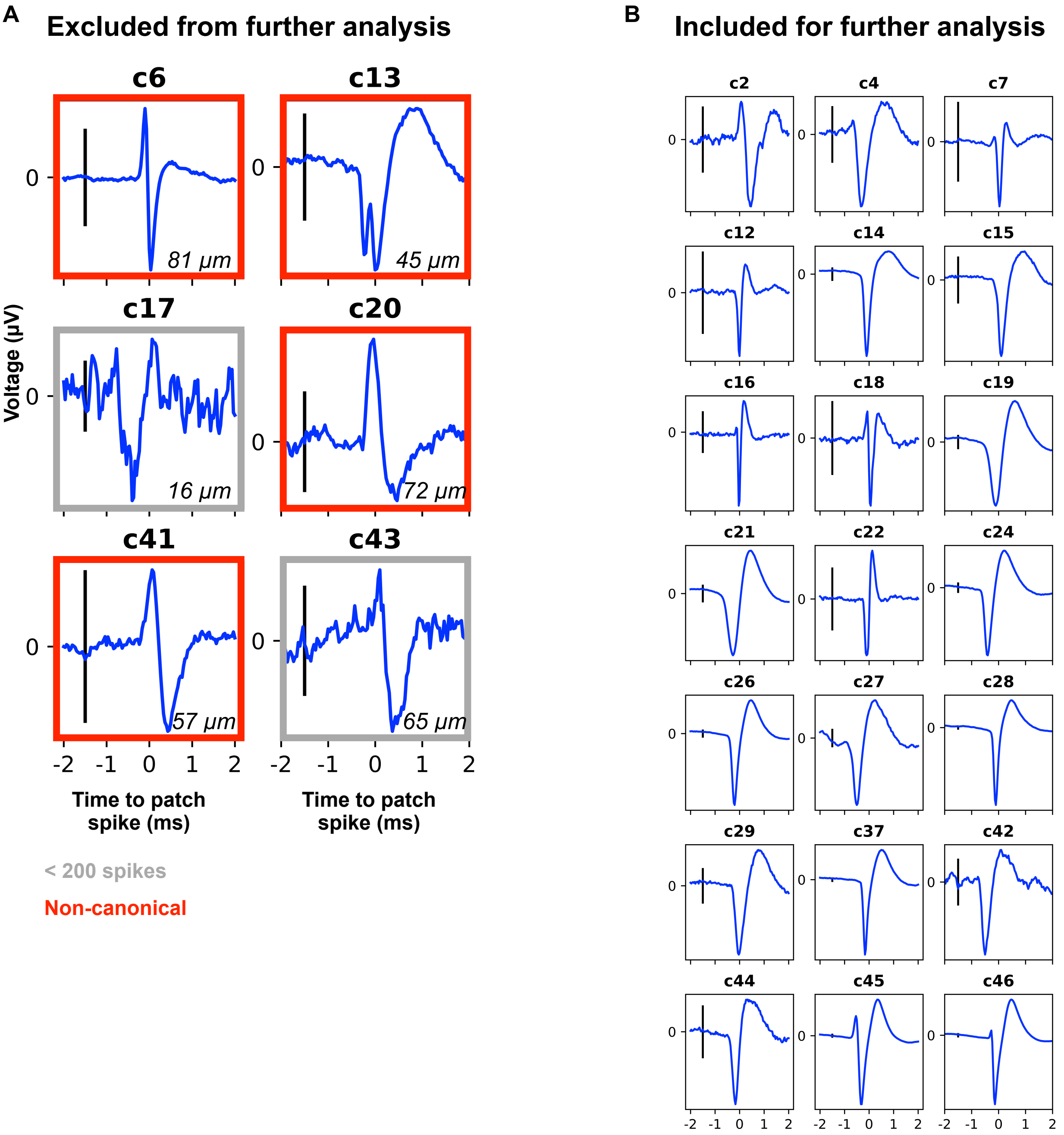


**Supplement to Figure 4-1.**

**A.** PSTA for the channel closest to the soma for six neurons which did not meet the inclusion criteria of firing sufficient number of spikes in patch-clamp (grey outline) or presenting a canonical spike waveform (red outline, see Materials and Methods). Scale bar is 10 $\mu$V. **B.** PSTA for the channel closest to the some for the 21 neurons meeting inclusion criteria. Scale bar is 10 $\mu$V.

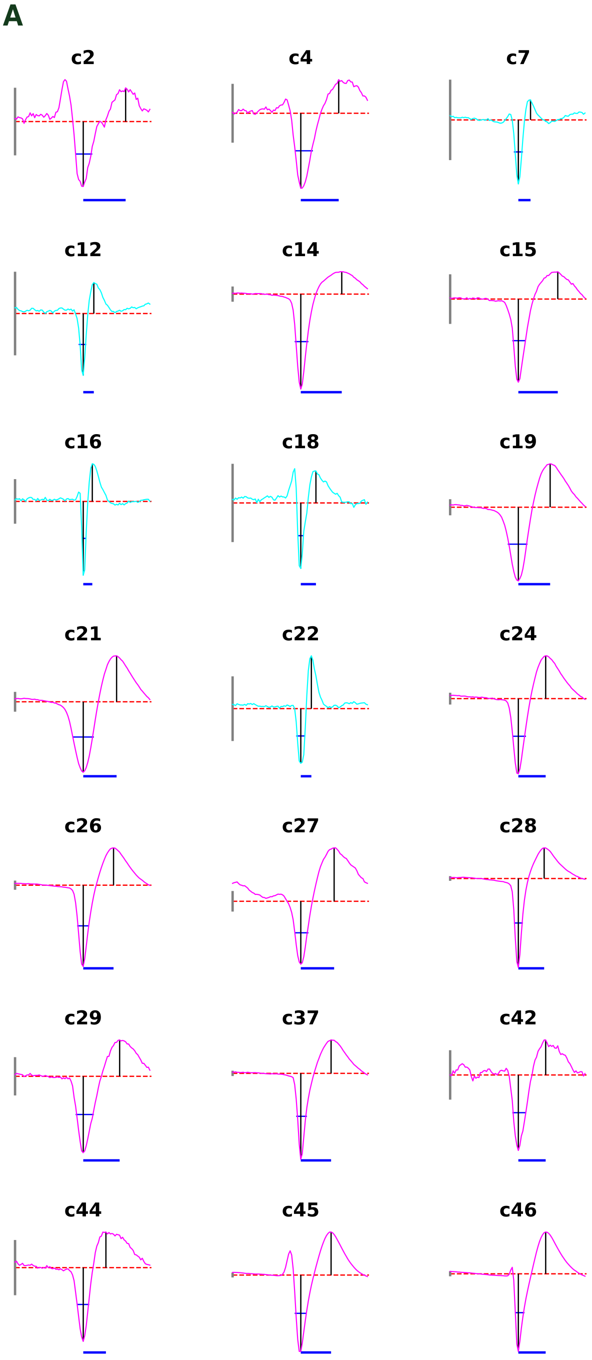


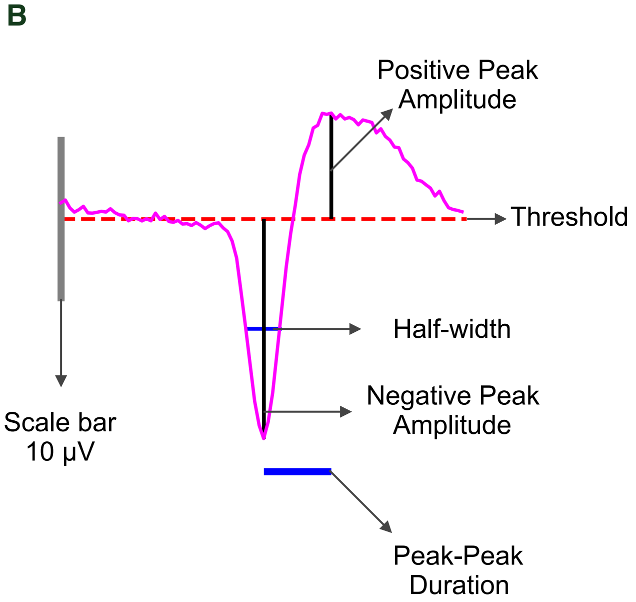


**Supplement to Figure 4-2.**

**A.** Measurement of spike features in PSTA waveforms for the 21 neurons passing the inclusion criteria for further analysis. Spike waveforms are coloured by putative cell type: PCs in magenta and INs in cyan.

**B.** Legend and example of feature extraction and measurement.

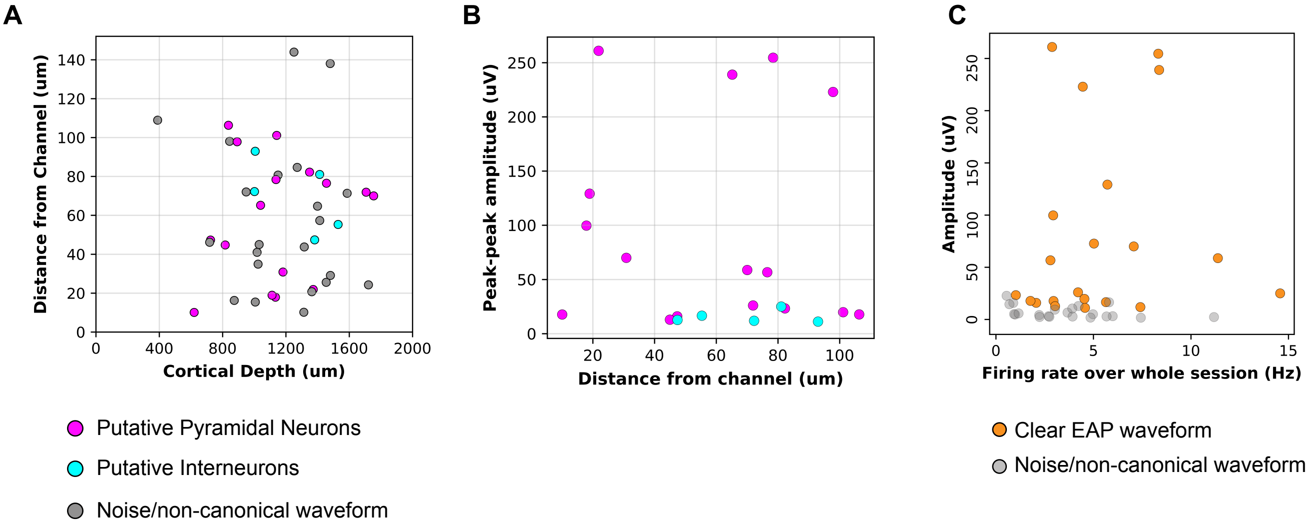


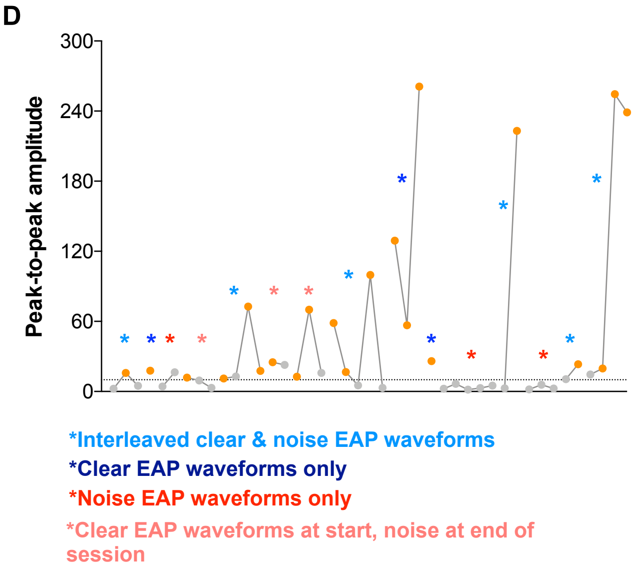


### Cells by Session Type

| | | | |
|---|---|---|---|
| 1 | c1, c2, c3 | 8 | c21, c22, c23, c24, c25 |
| 2 | c4 | 9 | c26, c27, c28 |
| 3 | c5, c6 | 10 | c29 |
| 4 | c7, c8, c10 | 11 | c31, c32, c33, c34, c35 |
| 5 | c12, c13, c14, c15 | 12 | c36, c37 |
| 6 | c16, c17 | 13 | c38, c39, c40 |
| 7 | c18, c19, c20 | 14 | c41, c42 |
| | | 15 | c43, c44, c45, c46 |

**Supplement to Figure 4-3.**

**A.** Distance from the closest extracellular channel versus cortical depth for every cell recorded. **B.** PSTA peak-peak amplitude versus distance in pyramidal cells (magenta) vs putative interneurons (cyan). **C.** PSTA amplitude versus firing rate averaged over the whole session for neurons with (orange) versus those without (grey) a clear EAP waveform after PSTA. **D.** Peak-peak amplitude for every neuron recorded ordered by session. Neurons from the same recording session are connected by a line. Cells in each session are detailed in a table (right). Session classification annotated by colour.
