## Supplement to Figure 5 for "Recording from the same neuron with high-density CMOS probes and patch-clamp: a ground-truth dataset and an experiment in collaboration"

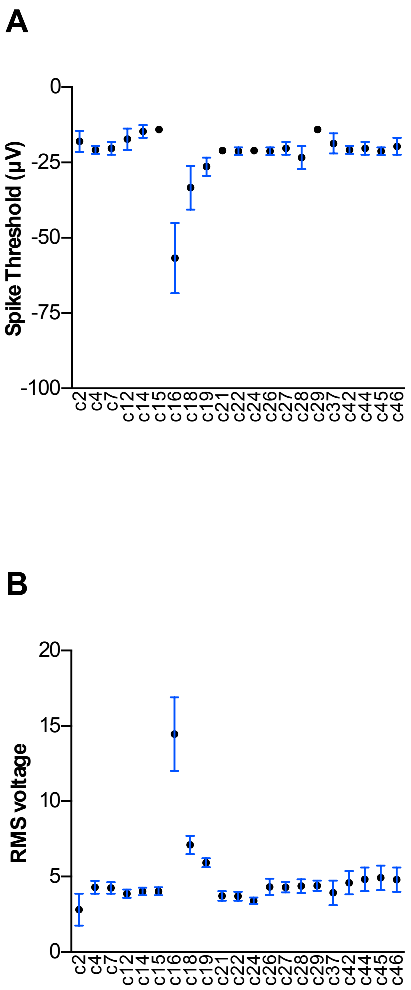


**Supplement to Figure 5. Extracellular detection of the patched neuron's action potentials.**

**A.** Spike threshold used per cell (mean ± SD for 30 channels). **B.** Root mean square voltage per cell (mean ± SD for 30 channels). **C.** PETHs between patch spike times and spike detections on the 30 probe channels closest to the soma for the subset of neurons with highest detectability, used for subsequent analysis. Legend as in Figure 4A.

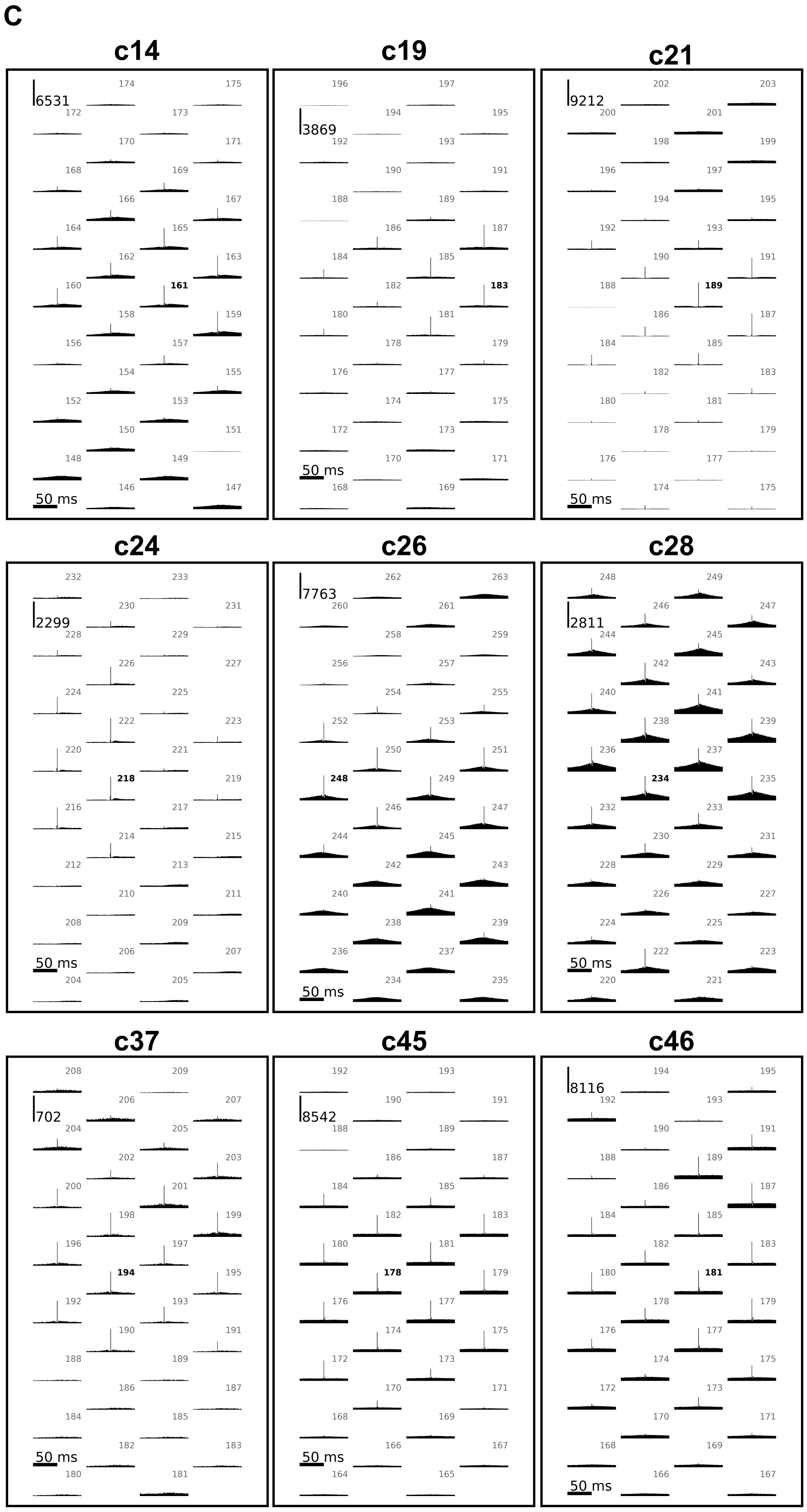
