## Supplement to Figure 6 for "Recording from the same neuron with high-density CMOS probes and patch-clamp: a ground-truth dataset and an experiment in collaboration"

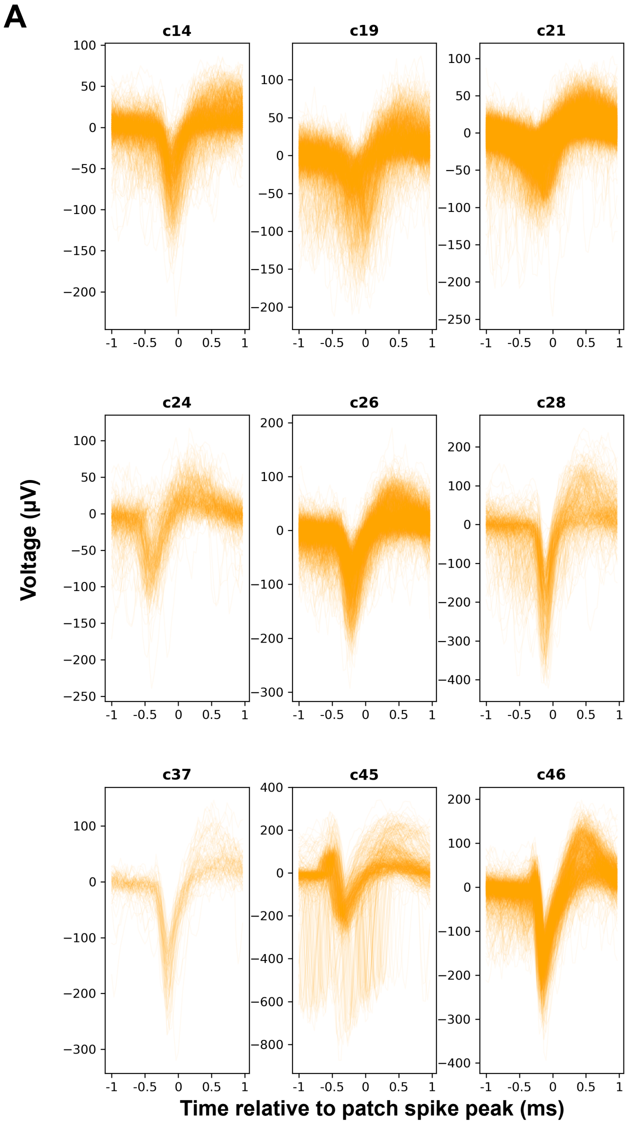


**Supplement to Figure 6-1. Trials excluded from reliability analysis.**

**A.** All spike traces excluded from reliability analysis. Temporal overlap with background units can be observed in each cell.

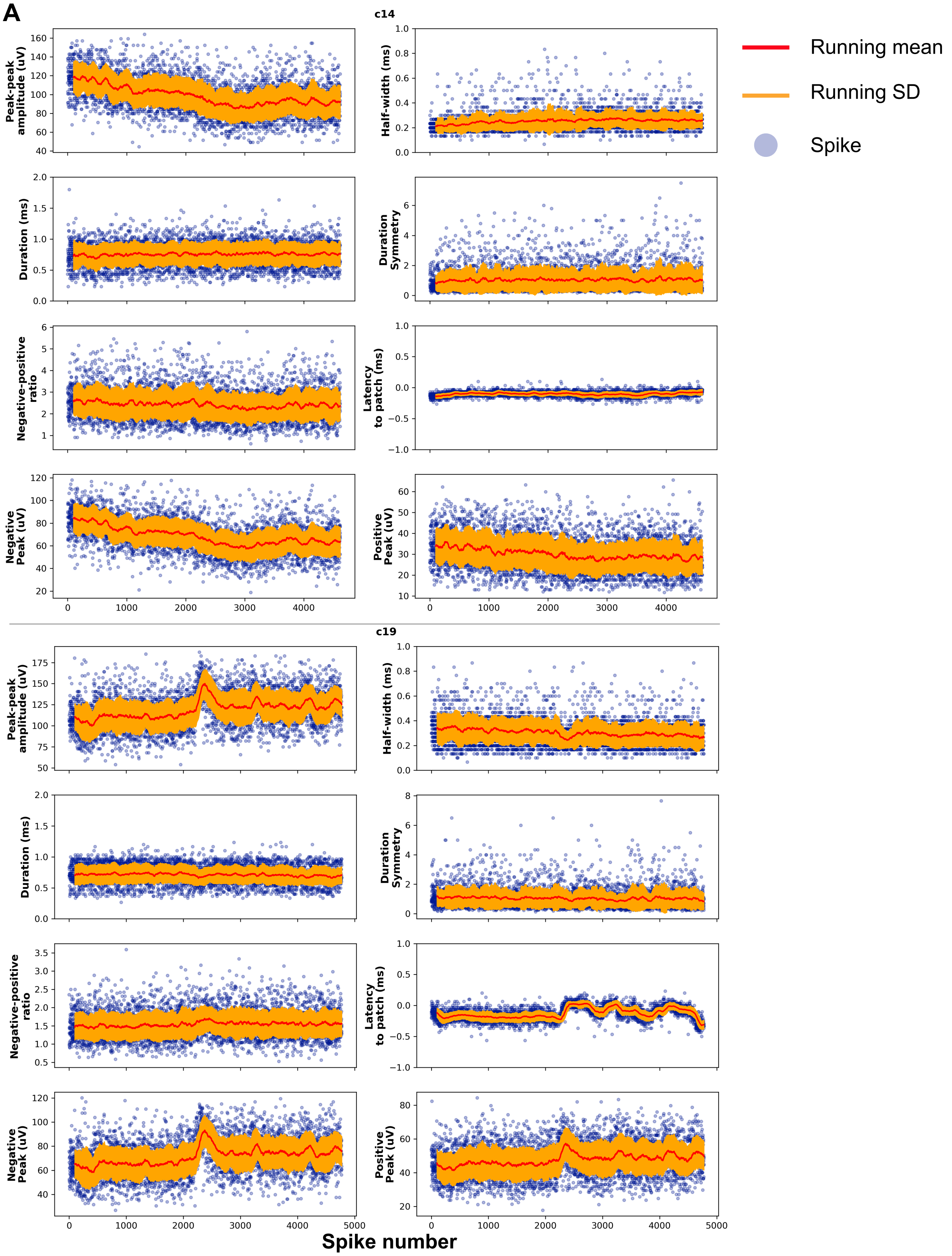


**Supplement to Figure 6-2. Variability of spike features over time.**

**A.** Running mean ± running SD (window of 100 spikes) and individual spikes for each of 8 spike features analysed over the length of each recording.

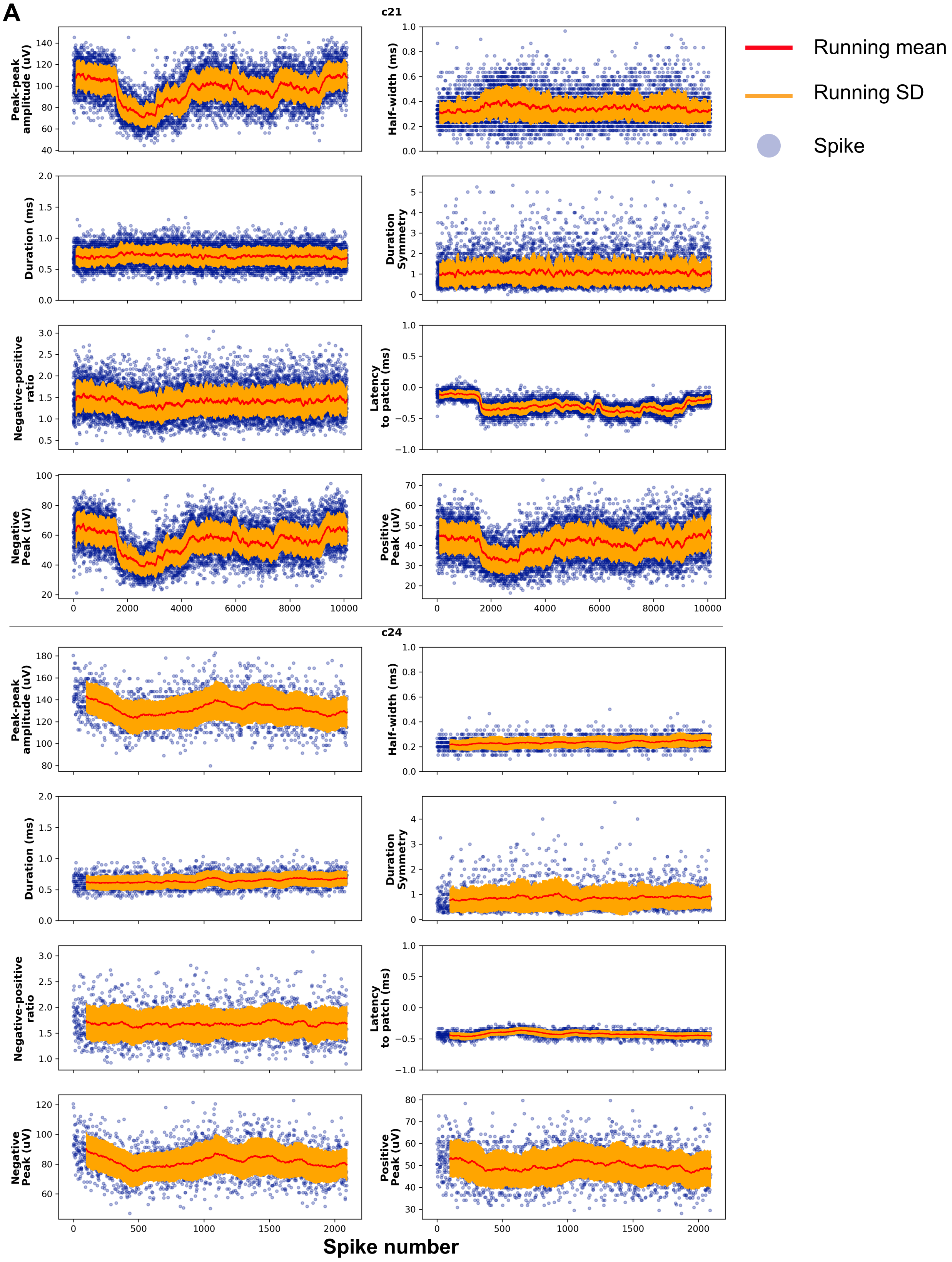


**Supplement to Figure 6-3. Variability of spike features over time.**

**A.** Running mean ± running SD (window of 100 spikes) and individual spikes for each of 8 spike features analysed over the length of each recording.

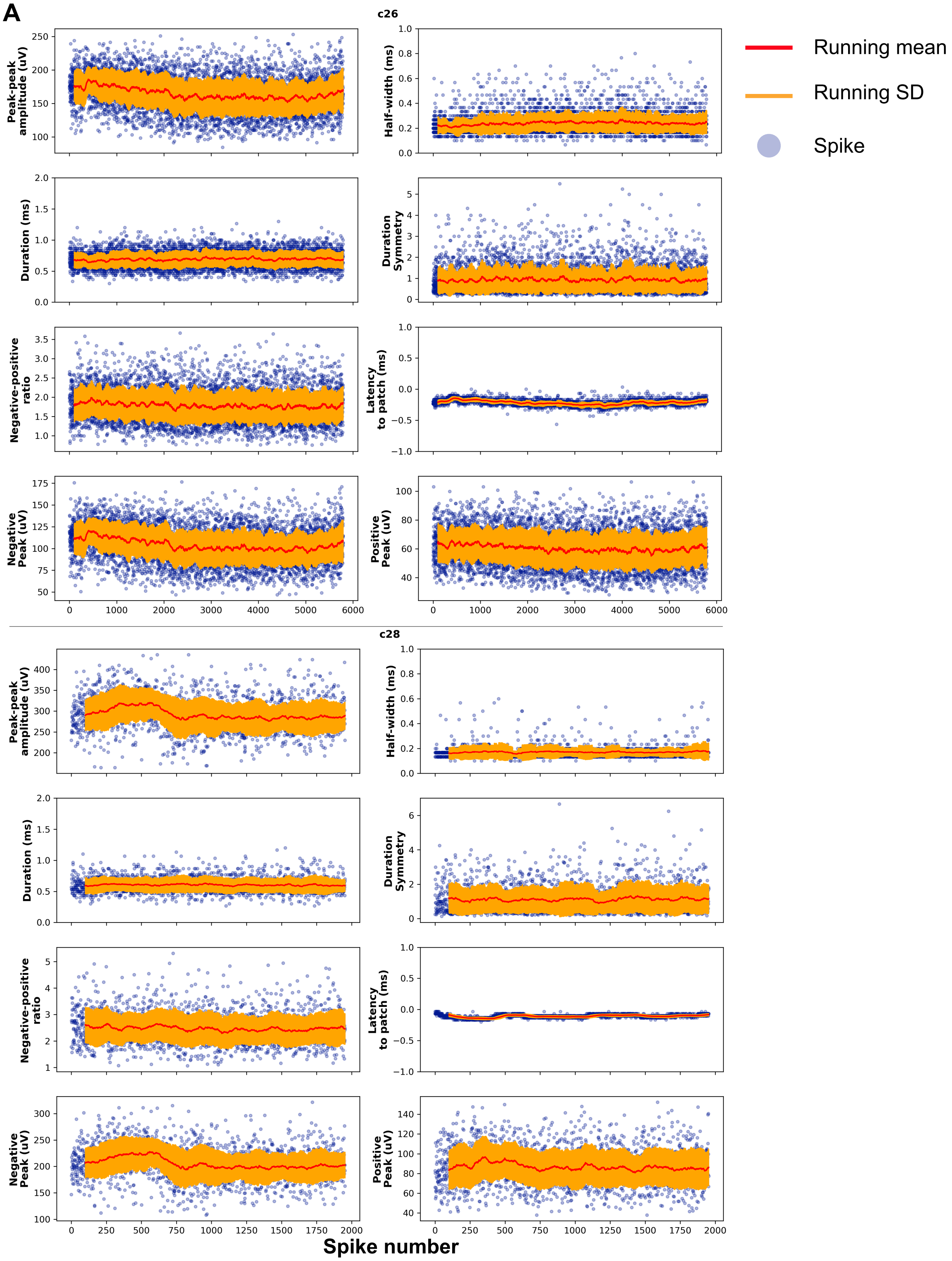


**Supplement to Figure 6-4. Variability of spike features over time.**

**A.** Running mean ± running SD (window of 100 spikes) and individual spikes for each of 8 spike features analysed over the length of each recording.

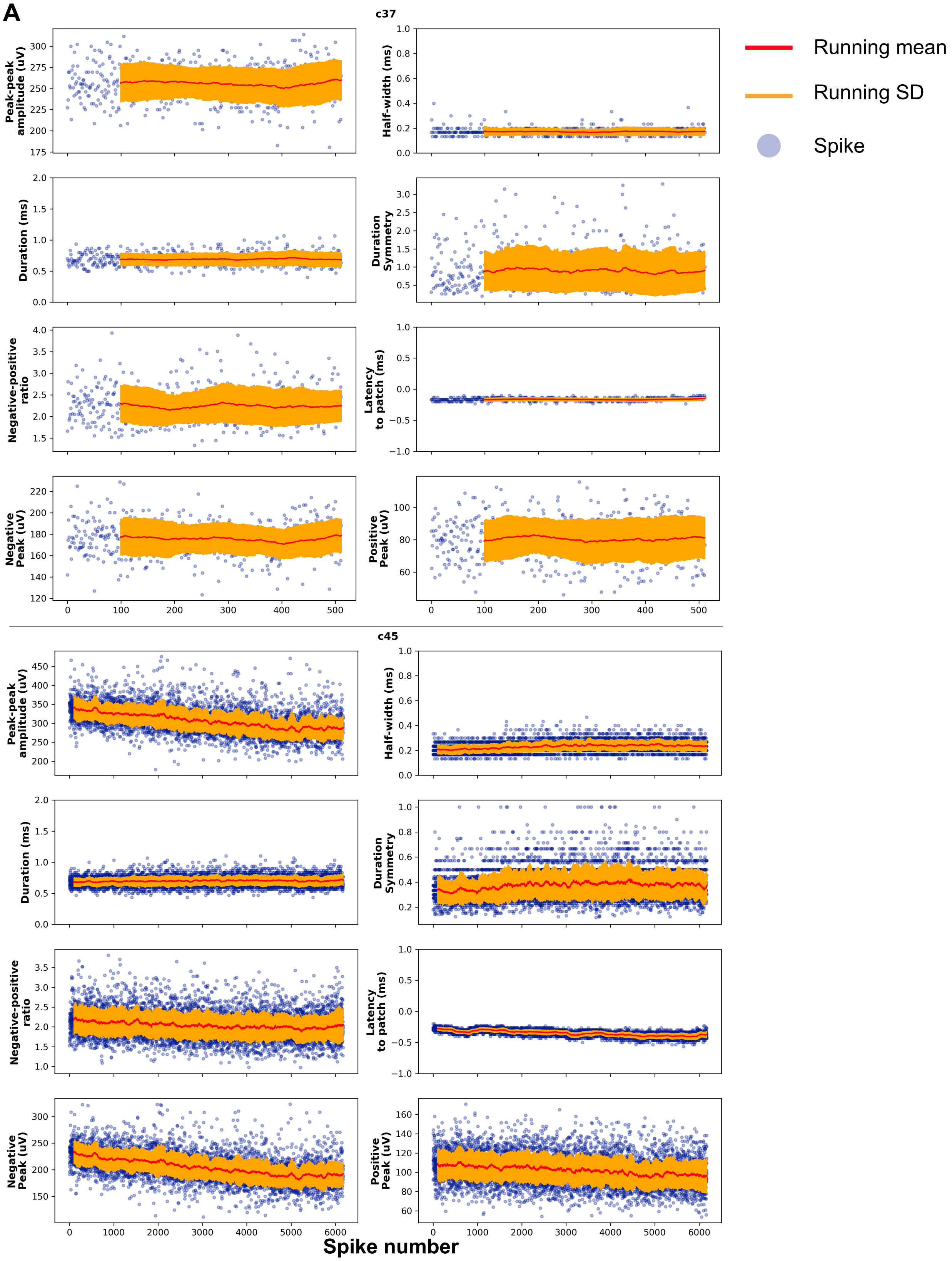


**Supplement to Figure 6-5. Variability of spike features over time.**

**A.** Running mean ± running SD (window of 100 spikes) and individual spikes for each of 8 spike features analysed over the length of each recording.

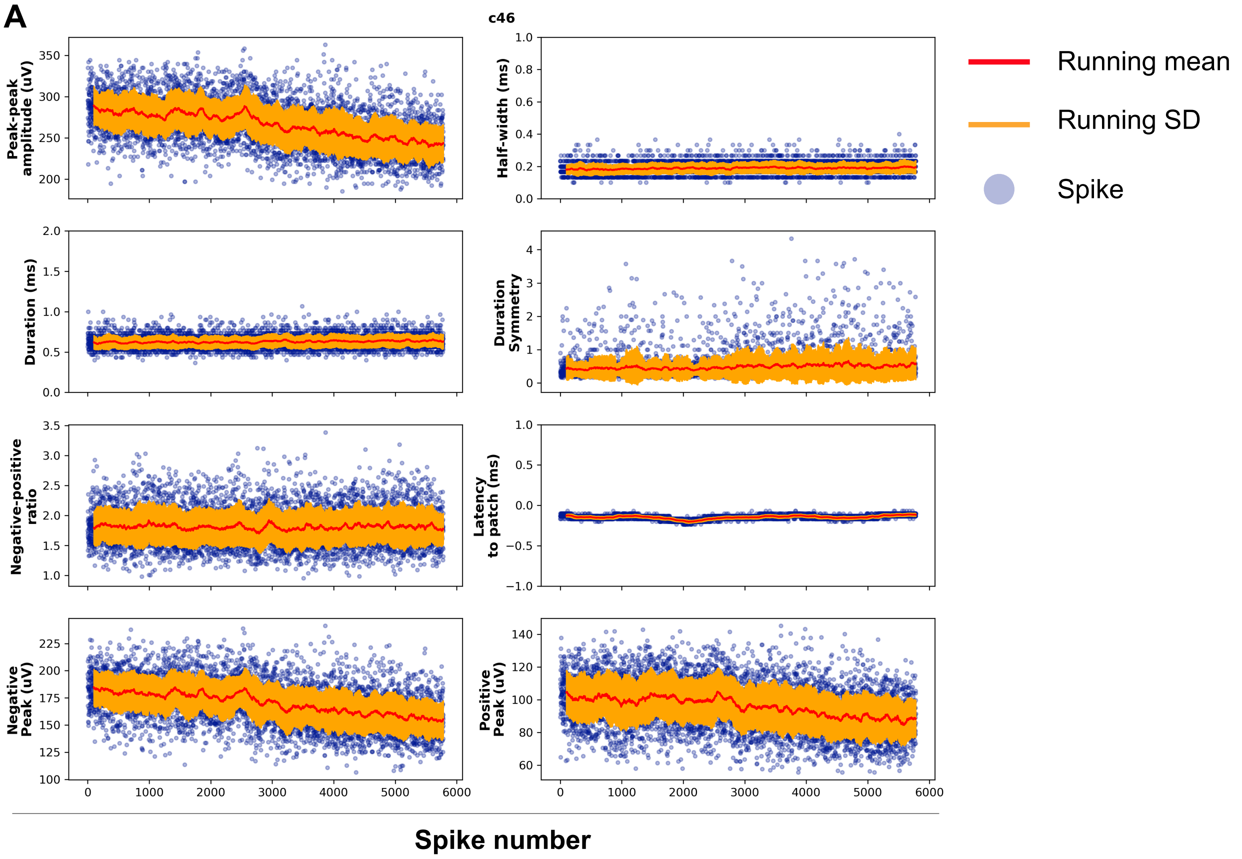


**Supplement to Figure 6-6. Variability of spike features over time.**

**A.** Running mean ± running SD (window of 100 spikes) and individual spikes for each of 8 spike features analysed over the length of each recording.
