## Supplement to Figure 7 for "Recording from the same neuron with high-density CMOS probes and patch-clamp: a ground-truth dataset and an experiment in collaboration"

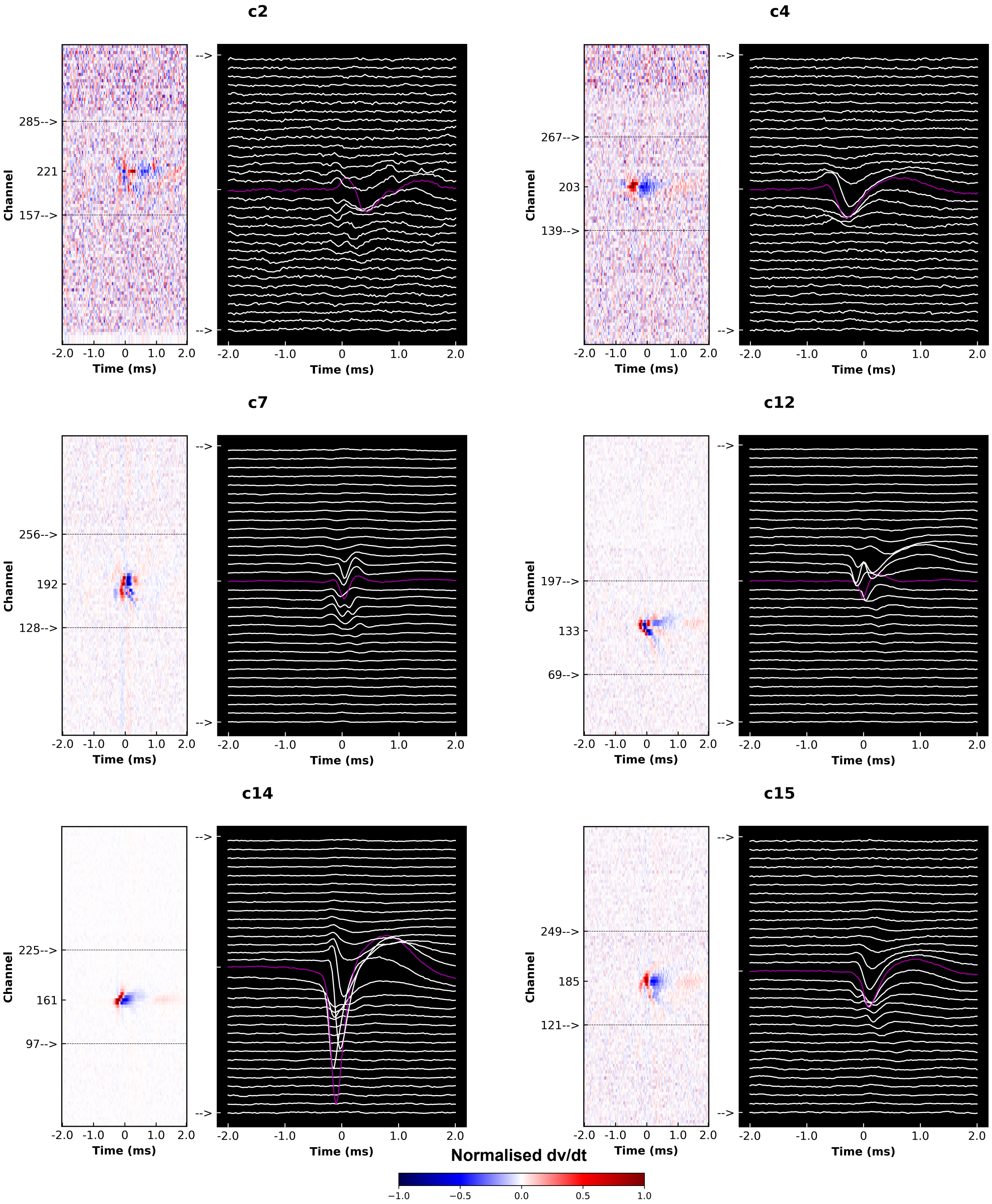


**Supplement to Figure 7-1. Spatiotemporal dynamics of extracellular action potential waveforms.**

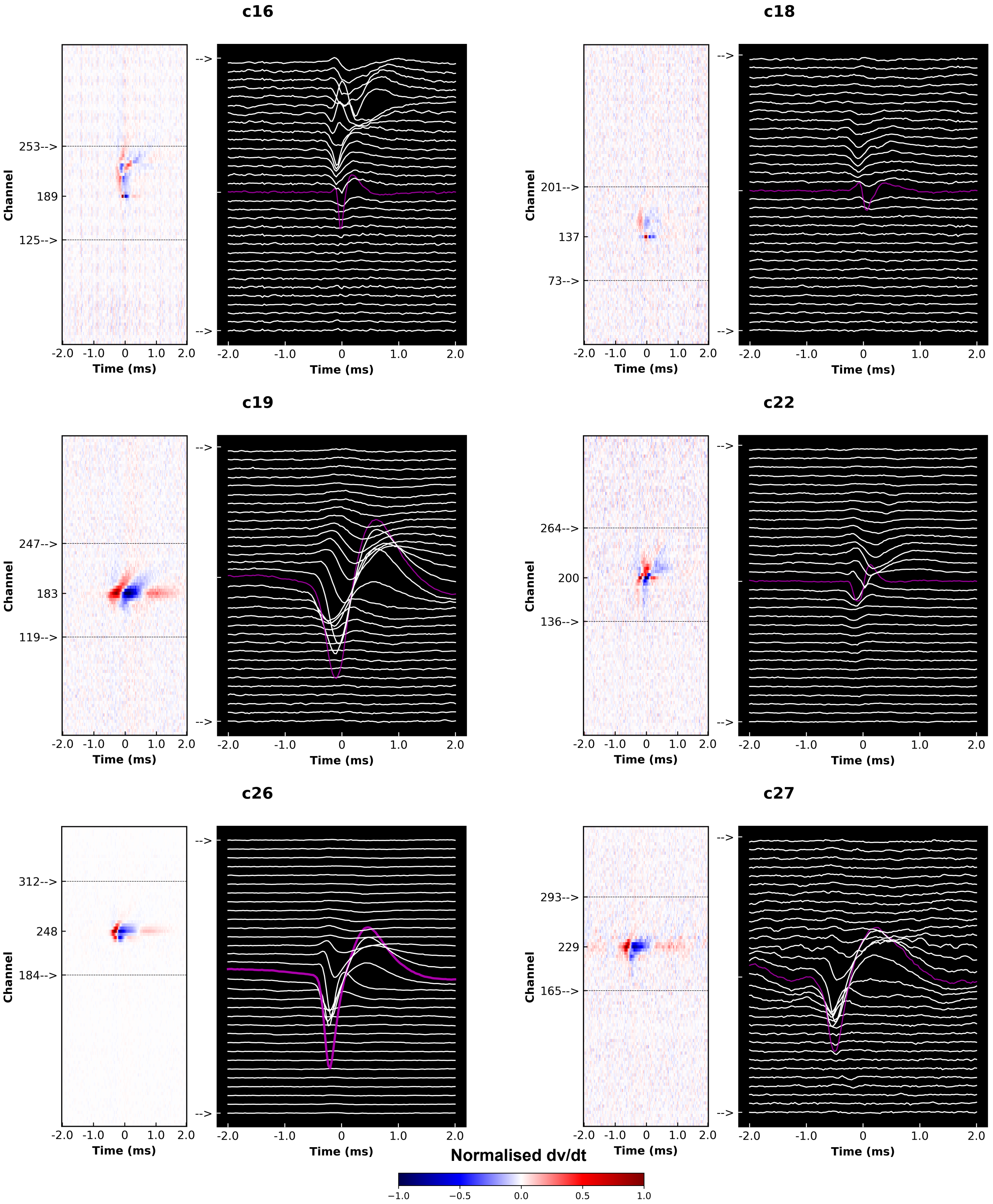


**Supplement to Figure 7-2. Spatiotemporal dynamics of extracellular action potential waveforms.**

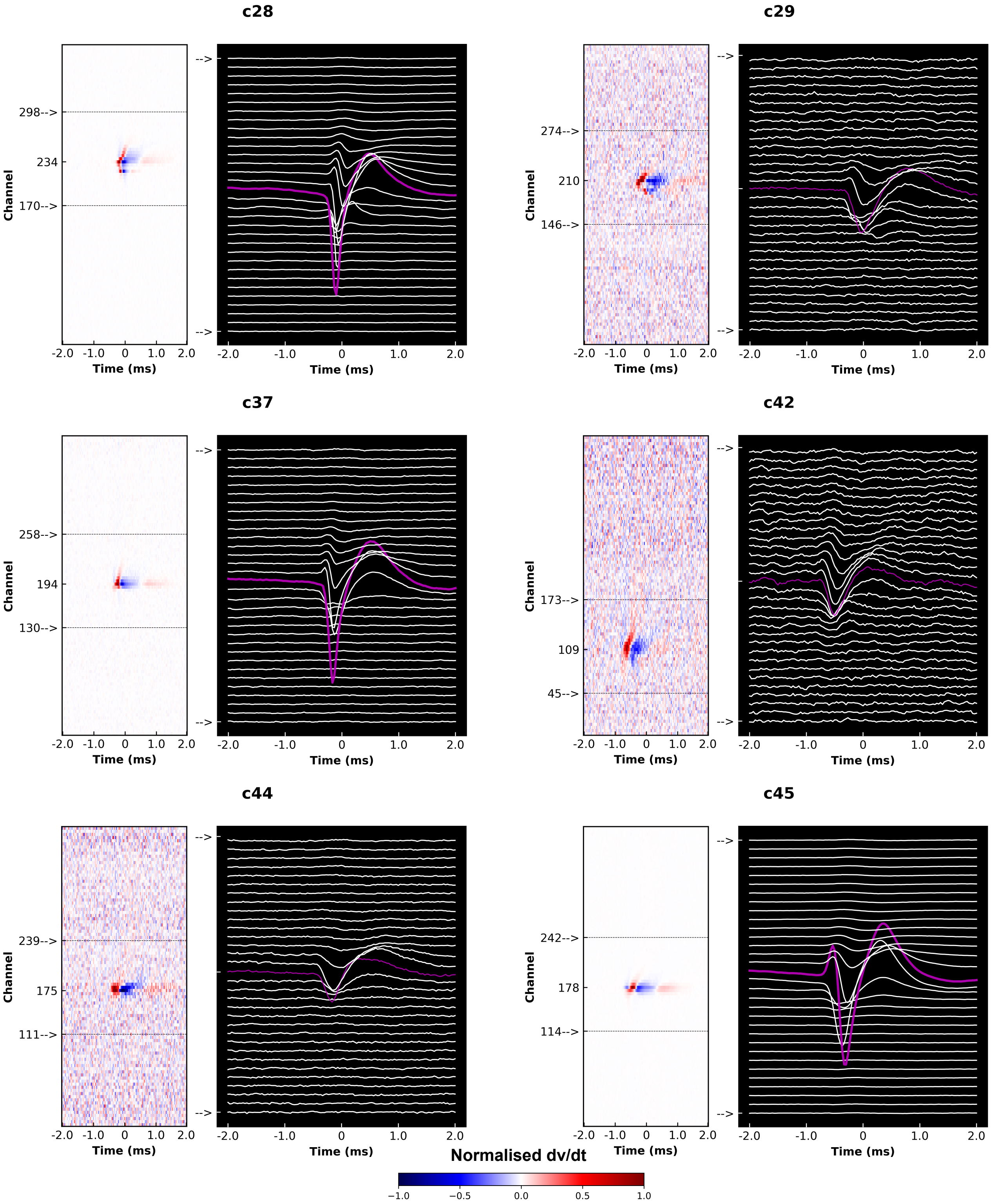


**Supplement to Figure 7-3. Spatiotemporal dynamics of extracellular action potential waveforms.**

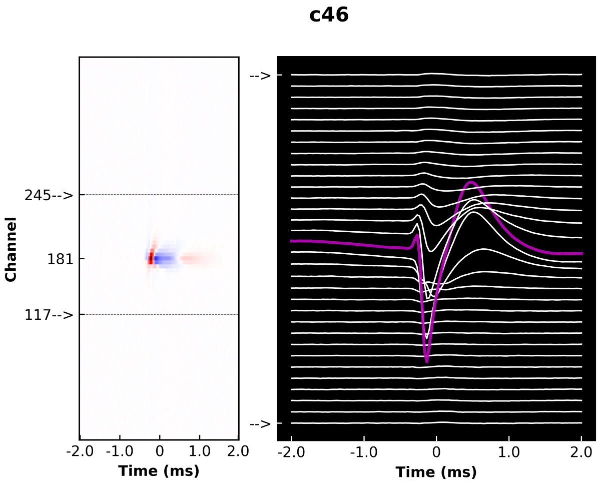


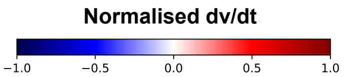


**Supplement to Figure 7-4. Spatiotemporal dynamics of extracellular action potential waveforms.**

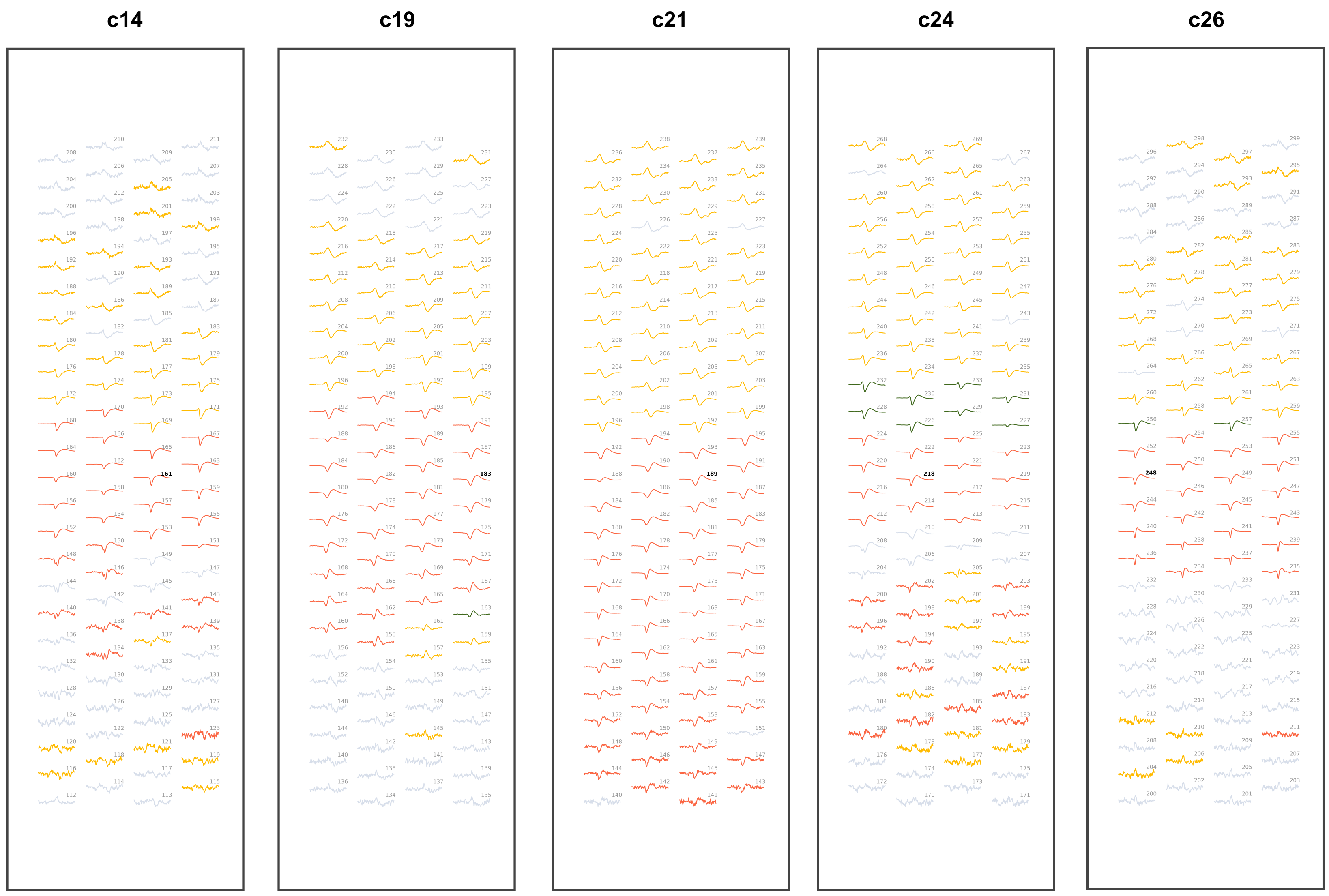


**Supplement to Figure 7-5. EAP waveform classification for 100 channels in the sample of 10 putative pyramidal cells analysed.**

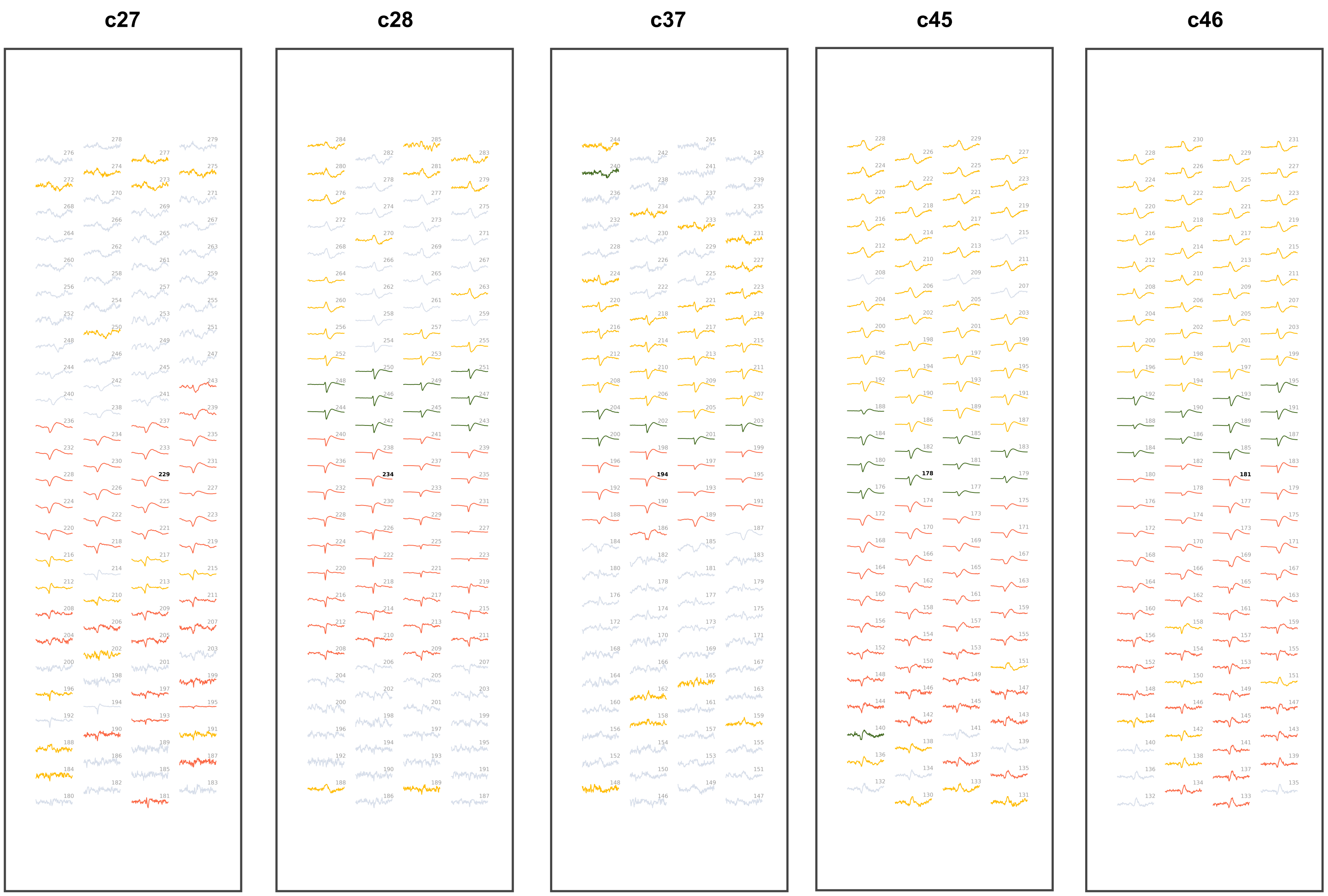


**Supplement to Figure 7-6. EAP waveform classification for 100 channels in the sample of 10 putative pyramidal cells analysed.**
